# Neuronal SKN-1B Modulates Nutritional Signalling Pathways and Mitochondrial Networks to Control Satiety

**DOI:** 10.1101/2020.07.21.213504

**Authors:** Nikolaos Tataridas-Pallas, Maximillian Thompson, Alexander Howard, Ian Brown, Marina Ezcurra, Ziyun Wu, Timo Keurten, Isabel Goncalves Silva, T. Keith Blackwell, Jennifer Tullet

**Author notes:** Joint first authors (Nikolaos Tataridas-Pallas, Maximillian Thompson).

## Abstract

The feeling of hunger or satiety results from integration of the sensory nervous system with other physiological and metabolic cues. This regulates food intake, maintains homeostasis and prevents disease. In *C. elegans*, chemosensory neurons sense food and relay information to the rest of the animal via hormones to control food-related behaviour and physiology. Here we identify a new component of this system, SKN-1B which acts as a central food-responsive node, ultimately controlling satiety and metabolic homeostasis. SKN-1B, an ortholog of mammalian NF-E2 related transcription factors (Nrfs), has previously been implicated with metabolism and respiration, because can mediate the increased lifespan incurred by dietary restriction. We show that actually SKN-1B is not essential for dietary restriction longevity and instead, controls a variety of food-related behaviours. It acts in two hypothalamus-like ASI neurons to sense food, communicate nutritional status to the organism, and control satiety and exploratory behaviours. This is achieved by SKN-1B modulating endocrine signalling pathways (IIS and TGF-β), and by promoting a robust mitochondrial network. Our data suggest a food-sensing and satiety role for mammalian Nrf proteins.

## Introduction

It is necessary for animals to correctly sense and adapt to food. Information on food cues is obtained via the sensory nervous system, integrated in the hypothalamus, and influences decisions about development, growth and behaviour (Bouret, 2017). These signals dictate appropriate food intake and regulate metabolic homeostasis, but are not well understood. In the nematode worm *C. elegans*, chemosensory neurons detect nutritional status, and relay this information to other tissues via hormones (Bargmann, 2006). These hormones activate downstream intracellular mechanisms including the insulin/IGF-1-like signalling (IIS) and transforming growth factor-β (TGF-β) pathways which act to switch behaviour between roaming (looking for and consuming food), dwelling (consuming food) and quiescence (a sleep-like state linked to satiety) depending on nutritional availability (Ben Arous et al., 2009; McCloskey et al., 2017; Skora et al., 2018; You et al., 2008). Adaptation to food cues also requires physiological changes, and mitochondrial networks are modulated to maximise energy output (Sebastián et al., 2017). Combined, these appropriate behavioural and physiological changes mean that food levels are correctly perceived, nutrient intake is regulated, and metabolic balance is maintained.

In mammals the NF-E2 related transcription factors (Nrfs) regulate a variety of processes. Nrf2 is known as a key, inducible, oxidative stress response factor but along with other Nrfs has also be implicated in proteostasis and metabolism (Blackwell et al., 2015). *C. elegans*, has only one sequence and functional Nrf orthologue, SKN-1, but its outputs are thought likely to be distributed between all the mammalian Nrfs (Blackwell et al., 2015). There are three *skn-1* isoforms (SKN-1A-C). SKN-1A and SKN-1C are expressed in the intestine and regulated, similarly to the Nrfs, at the level of cellular localisation (Lehrbach and Ruvkun, 2016; Tullet et al., 2008). In contrast, SKN-1B is expressed in two chemosensory neurons, the ASIs, which are thought to act as the worm’s hypothalamus, and is constitutively nuclear (Bargmann, 2006; Bishop and Guarente, 2007; Tullet et al., 2008). SKN-1B has been of particular interest with respect to metabolism and respiration, because its action in ASI can mediate the increased lifespan incurred by dietary restriction (DR) (Bishop and Guarente, 2007).

We further tested the role of SKN-1B in DR mediated longevity but found it to be non-essential. Instead, we identify SKN-1B to be deeply ingrained in food-detection and food-related behavioural responses. Specifically, we find that SKN-1B: regulates satiety in response to fasting and re-feeding; promotes exploration in fed conditions; and controls appropriate responses to fasting. Our data suggest that SKN-1B controls food-related behaviour both via modulating the key signalling pathways (TGF-β and insulin signalling), and physiologically through the control of mitochondrial networks. This places SKN-1B at the heart of food-responsive signalling pathways, where it acts to regulate satiety and control metabolic homeostasis. Our data suggest the possibility that Nrfs act to regulate food-sensing and satiety in humans.

## Results

### SKN-1B contributes to DR longevity, but is not necessarily essential

SKN-1 is a well characterised promoter of longevity: Mutants lacking all *skn-1* isoforms are short lived and mild overexpression of SKN-1 extends lifespan (Blackwell et al., 2015; Lehrbach and Ruvkun, 2019). In particular, expression of *skn-1b* in the ASI neurons can mediate the extension in lifespan incurred by a food dilution DR protocol, suggesting that SKN-1B might be a general and essential mediator of DR (Bishop and Guarente, 2007). Multiple *C. elegans* DR protocols exist, some of which have different underlying genetic requirements (Kapahi et al., 2017), so we explored the specific requirement for *skn-1b* for these other forms of DR. The weaker, ~20% lifespan extension observed in *eat-2* mutants required *skn-1b* (Figure 1A and Table S1. However, an alternative food dilution protocol that extends WT lifespan more dramatically ~40-60%, and is dependent on *skn-1* (Moroz et al., 2014), proved independent of *skn-1b* (Figure 1B and Table S2). We conclude that although *skn-1b* contributes to DR mediated longevity under some conditions, it is not necessarily essential. Like DR, rIIS extends lifespan in many species and *skn-1* is known to be an important mediator of this (Ewald et al., 2014; Tullet et al., 2008). However, *skn-1b* was not required for the long life of *daf-2* mutants, suggesting either redundancy among isoforms or a requirement for other isoforms in particular (Figure S1A-F, Figure S2, Tables S3 and S4). Neither did we observe any requirement of *skn-1b* for WT lifespan (Figures S1 and S2, Tables S3 and S4). In summary, *skn-1b* does not contribute to longevity under normal or rIIS conditions, but does contribute to the lifespan incurred by specific DR conditions.

**Figure 1.**
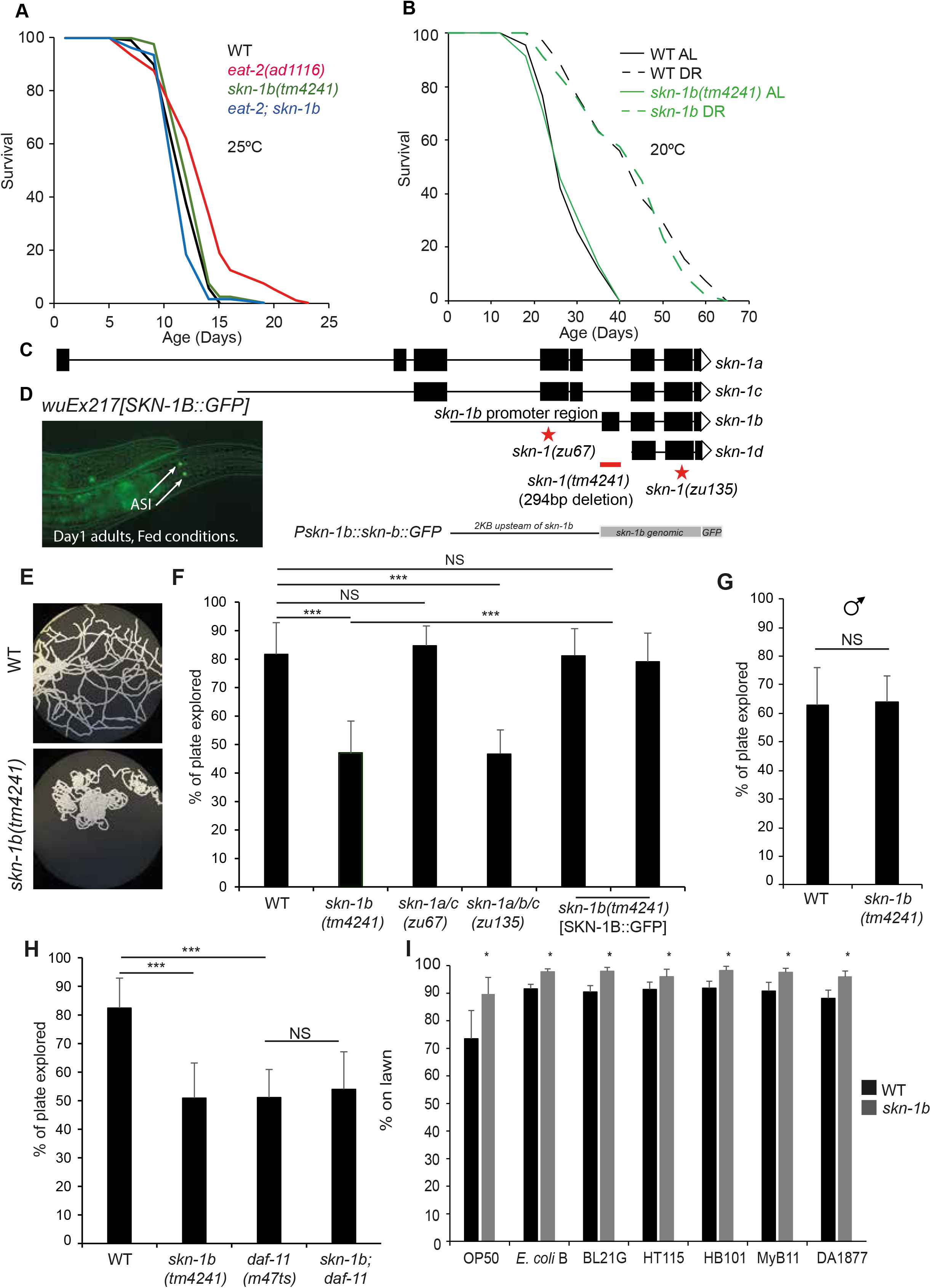
*skn-1b* is required for exploratory behaviour, but is not essential for DR longevity. **A)** Effect of *skn-1b* on *eat-2* lifespan. **B)** Survival of WT and *skn-1b* mutants in response to bacterial dilution as in (Moroz et al., 2014). For **A and B**: Representative experiments shown, individual trials summarised with Log-Rank analysis in Tables S1 and S2. These DR protocols did not alter SKN-1B::GFP levels (Figures S7B and S7C). We use *eat-2* as a DR longevity model as suggested (Lakowski and Hekimi, 1998), but recent work shows that *eat-2* longevity also derives from reduced pathogenesis (Kumar et al., 2019; Zhao et al., 2017). *C. elegans* derives nutrients from the bacteria and removing pathogenic components of bacteria will undoubtedly alter its nutritional profile but separating the two is challenging. **C)** Genetic locus of *skn-1* with isoforms, mutants and SKN-B::GFP specific transgene. *skn-1b* mRNA is not detectable in *tm4241* mutants but *skn-1a* and *c* mRNA levels are unchanged implying that this allele is *skn-1b* specific (Figure S3A)*. skn-1b* mutants have normal brood sizes (Figure S3B-E). *skn-1(zu67)* and *skn-1(zu135)* encode point mutations leading to early stop codons and transcript degeneration via non-sense mediated decay. All *skn-1* isoforms have the same binding site, and all Nrfs can bind the same sequence, suggesting the likelihood of overlapping targets. **D)** *wuEx217* SKN-1B::GFP is expressed in ASI neurons. The SKN-1B::GFP translational reporter confirmed that SKN-1B::GFP can be expressed independently from other SKN-1 isoforms, that *skn-1b* is expressed solely in the ASIs. This expression pattern was confirmed with an endogenous Scarlet::SKN-1B reporter (Figure S3F). SKN-1B::GFP expression varies at different ages (Figure S3G). ASIs confirmed by DiI staining and SKN-1B::GFP was rarely observed in additional neurons (Figure S3H). **E)** Agar plates showing exploration of a single worm over 16hrs. Assay and controls shown (Figure S4). **F-H)** Quantification of exploration. Mean plate coverage of n>11 worms per group ± st. dev. One representative experiment of 3 biological replicates shown. In **H)** SKN-1B was rescued using the *ukcEx15* and *ukcEx16* transgenes. In **G)** a 2hr period was used to allow quantification of hyperactive male exploration. **I)** Quantification of worms on different small bacterial lawns. Assay (Figure S5A). Each bar represents a mean of 3 biological replicates ± st. dev. For **F-I)** Two-tailed *t*-test \**p*<0.05, \*\**p*< 0.001, \*\*\**p*<0.0001, NS not significant.

### *skn-1b* acts to promote food-related exploratory behaviour

Sensory input via the ASIs affects *C. elegans’* three main food-related locomotory behaviours (roaming, dwelling and quiescence) (Bargmann, 2006; Trojanowski and Raizen, 2016). Given that SKN-1B is implicated in DR longevity we explored the role of *skn-1b* in behaviour using a *skn-1b*-specific allele *(tm4241)* (Figure 1C and Figure S3A-E). To gain an overview of food-related behavioural patterns, we quantified the ability of *skn-1b* mutants to “explore” a continuous bacterial lawn during a 16hr period compared to WT, an assay shown to correlate with classical roaming and dwelling assays (Flavell et al., 2013) (Figure S4A). During this period, WTs explored ~80% of the lawn, but *skn-1b* mutants only explored ~45% suggesting that *skn-1b* mutants’ exhibit reduced exploratory behaviour (Figures 1E and 1F). We observed similar behaviour in *skn-1(zu135)* mutants which lack all *skn-1* isoforms, but not in *skn-1(zu67)* mutants which are thought to lack only *skn-1a* and *c* (Figures 1C and 1F). Furthermore, rescuing *skn-1b* mutants with a SKN-1B::GFP specific transgene, which drives *skn-1b* expression from its own promoter specifically in the ASIs, fully restored exploratory behaviour to WT levels (Figures 1D and 1F, Figure S3F).

As some *skn-1* isoforms are important for normal embryogenesis (Bowerman et al., 1993), it is possible that the *skn-1b* requirement for normal exploration could be due to disrupted ASI development. However, *skn-1* RNAi from the post-embryonic L1 or L4 stage was sufficient to decrease exploration, indicating that this phenotype is not due to a *skn-1b*-related embryonic development defect (Figure S4C-E). *skn-1b* mutants also performed as well as WT in an assay of thrashing behaviour indicating that their movement was not generally impaired (Figure S4F). We also explored behavioural differences in male *C. elegans* that have evolved to balance the competing needs of reproduction *versus* foraging. For instance, in the absence of hermaphrodites, males increase exploratory behaviour to search for mates (Barrios et al., 2008; Lipton, 2004). However, we found that both WT and *skn-1b* males explored to the same hyperactive degree (Figure 1G). Thus, *skn-1b* promotion of motility appears to support foraging rather than mate location. Together, we conclude that adult expression of *skn-1b* in ASIs contributes to normal exploratory behaviour.

The ASI neurons consist of cell bodies that reside anterior to the large bulb of the pharynx, with projections reaching forward to the amphid openings (the worm’s nose) (Bargmann, 2006). At the amphid openings, the ASIs express transmembrane receptor-type guanylate cyclases such as *daf-11* that relay environmental cues to the cell body (Birnby et al., 2000). *daf-11* mutants have sensory defects and fail to chemotax towards a number of attractants including NaCl and diacyl as well as being required for normal dauer entry and exit (Birnby et al., 2000). To explore the relationship between *skn-1b* and *daf-11* we tested their epistatic relationship in relation to behaviour. Similarly to *skn-1b* mutants, we observed an exploratory defect in *daf-11* mutants (Figure 1H) and notably, a *skn-1b; daf-11* double mutant did not exhibit a greater reduction in exploration (Figure 1H). The lack of an additive effect of these two mutations suggests that *daf-11* and *skn-1b* act in the same pathway to influence behaviour.

In exploration assays *C. elegans* are cultured on a continuous lawn of *E. coli.* As *skn-1b* mutants explore less, we reasoned that they may spend less time away from food than WTs. To test this, we provided the worms with a small lawn of OP50 bacteria in the centre of an otherwise empty plate, and counted the number of worms on and off the bacteria (Figure S5A). Whilst at any given time approximately 25% of WT worms are off a standard OP50 lawn, at the same point all *skn-1b* mutants remained on the lawn (Figure 1I). Similar mild avoidance of lawns in WT but not *skn-1b* mutants was seen for other bacteria, including another four *E. coli* strains, *Comamonas aquatica* and a *Pseudomonas sp.* (Figure 1I). However, when WT worms are fed *B. subtilis* the proportion on the lawn increases compared to OP50 whereas that of *skn-1b* mutants remains the same (Figure S5B). Similarly, no differences in lawn avoidance were seen on *E. coli* W3110 or MG1655 (Figure S5B). As almost all *skn-1b* mutants are present on an OP50 lawn, it implies that while WT worms adapt to preferentially consume some foods, *skn-1b* mutants do not. We also tested whether *skn-1b* might contribute to a pathogen avoidance response and examined food avoidance behaviour of WT and *skn-1b* mutants fed pathogenic *Pseudomonas aeruginosa.* However, both WT and *skn-1b* mutants avoided the pathogen to a similar extent indicating that *skn-1b* is not involved in pathogen avoidance behaviour (Figure S5C). Overall, this indicates that *skn-1b* acts to sense food types rather than pathogenicity and subsequently controls behaviour.

### *skn-1b* regulates behaviour in response to fasting

Exploration allows worms to seek and locate food (Ben Arous et al., 2009). When re-fed after a period of fasting, exploration is reduced and worms ‘dwell’ to increase their food consumption and refuel their energy stores, they then enter satiety quiescence (Ben Arous et al., 2009; You et al., 2008). These responses are regulated by the ASIs and hormones, so we investigated the contribution of *skn-1b.* We fasted WT and *skn-1b* mutants for 1hr and quantified their behaviour upon re-feeding. Whilst WT worms exhibited the expected reduction in exploration under these conditions, *skn-1b* mutants did not (Figure 2A). We also fasted WT and *skn-1b* mutants for 16hrs, and examined their exploration following re-feeding. We found that while this more extreme fasting protocol caused a marked decrease in WT activity compared to 1hr fasting, it had no effect on *skn-1b* mutants (Figures 2A and 2B). This demonstrates that *skn-1b* is required for behavioural control in response to fasting and re-feeding.

**Figure 2.**
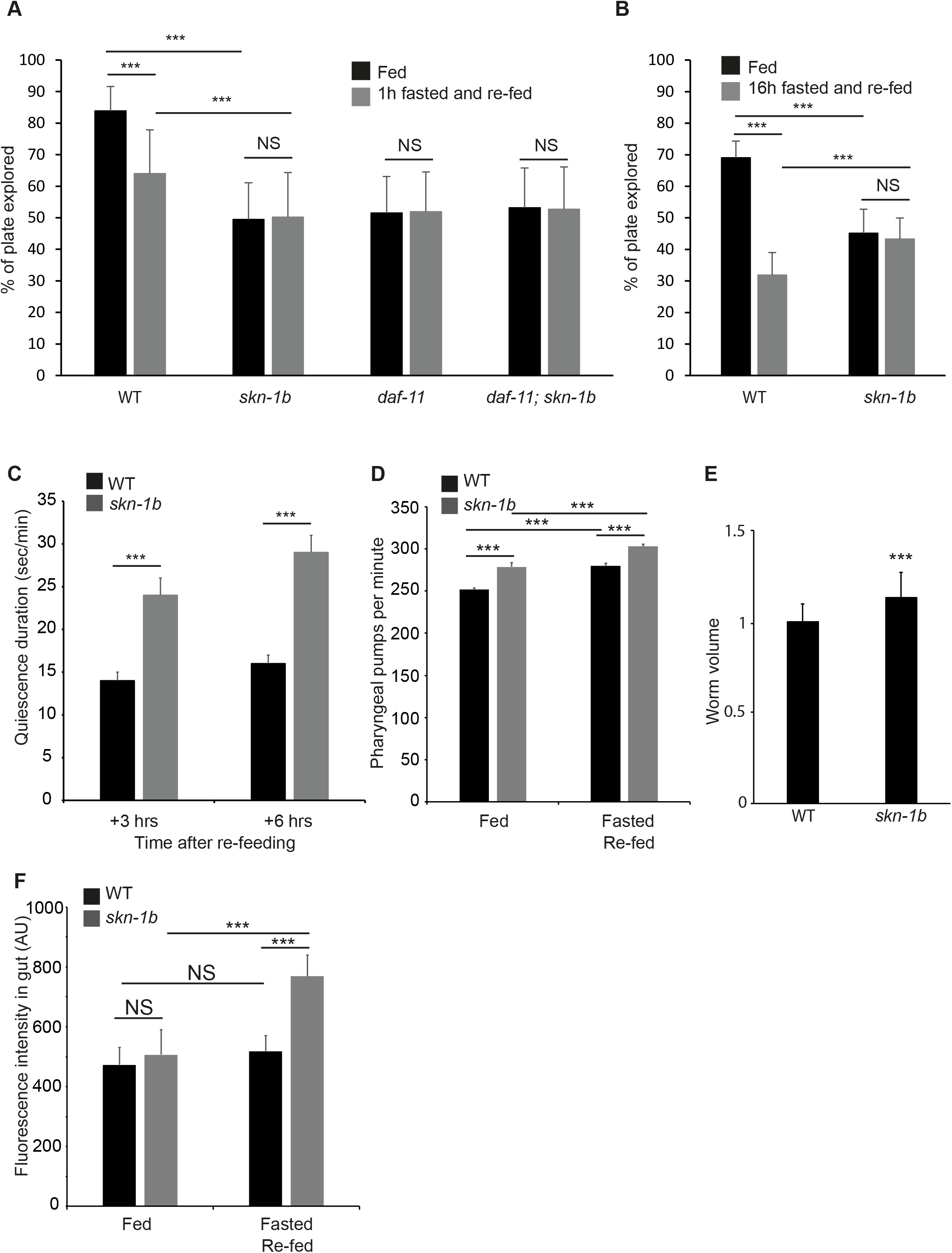
SKN-1B regulates satiety quiescence. **A)** Quantification of exploration in fed *vs* fasted/re-fed conditions, worms fasted for 1hr. Mean plate coverage of n>35 individual worms per group ± st. dev., 3 combined experiments shown. **B)** Quantification of exploration in fed *vs* fasted/re-fed conditions, worms fasted for 16hrs. Mean plate coverage of n>7 worms per group ± st. dev., one representative experiment of 3 trials shown. **C)** Time spent in quiescence after fasting/re-feeding. Each bar represents a mean of 3 biological replicates ± SEM, n>40 worms per group. **D)** Pharyngeal pumping rate. Each bar represents a mean of 3 biological replicates, ± st. dev., n=7 worms per group. **E)** Worm volume. Each bar represents a mean of 3 biological replicates, ± st. dev., n>63 worms per group. **F)** Effect of *skn-1b* on intake of fluorescently labelled OP50. Each bar represents a mean of 3 biological replicates, ± st. dev., n>42 worms per group. For **A-F**) Two-tailed *t*-test \**p*<0.05, \*\**p*< 0.001, \*\*\**p*<0.0001, NS not significant.

As *daf-11* mutants also exhibit decreased exploration, (Figures 1H and 2A), we tested whether *daf-11* was also required for behavioural changes in response to fasting and re-feeding. We found that *daf-11* worms, like *skn-1b* mutants, do not respond to fasting and re-feeding, and that the combined effect of *daf-11; skn-1b* mutation was non-additive (Figure 2A). That both DAF-11 and SKN-1B are required for worms to modulate their exploratory behaviour in response to fasting and re-feeding provides further evidence that these two proteins act in concert.

The decreased exploration observed in response to fasting and re-feeding can be attributed either to increased time spent dwelling, or in quiescence (Ben Arous et al., 2009; You et al., 2008). When dwelling, pharyngeal pumping is normal and *C. elegans* makes minimal back and forward sinusoidal movement (Bargmann, 2006). In contrast, quiescent worms do not pump or move at all (McCloskey et al., 2017; You et al., 2008). As *skn-1b* mutant’s exhibit reduced exploration we asked whether they also differ in these other behaviours. After fasting, WT worms are quiescent for a longer period during the 3-6hrs after re-feeding, making this the best time to measure satiety quiescence (McCloskey et al., 2017; You et al., 2008). We found that at both 3 and 6hrs after re-feeding, *skn-1b* mutants spent longer in a quiescent state than WT worms (Figure 2C). Similar numbers of WT and *skn-1b* mutants quiesce under these conditions (Figure S6A). Together, this suggests that SKN-1B acts to suppress satiety-induced quiescence promoting exit from, but not entry into quiescence. Although these assays do not measure quiescence in fed conditions, taken together with the inability of *skn-1b* mutants to modify their behaviour in response to fasting and re-feeding (Figure 2A and 2B), their results imply that, the reduced exploration in *skn-1b* mutants is due to increased time spent in quiescence.

Quiescence is linked to satiety in mammals, and quiescent *C. elegans* do not pump food into their gut, so these data could imply that *skn-1b* mutants eat less compared to WT. In addition to the time spent on food, the amount of food that a worm eats can be determined by the efficiency and rate of pharyngeal pumping and the amount of time that it spends pumping (You et al., 2008). To test this, we compared pumping rate in fed WT and *skn-1b* mutants and observed a modest but statistically significant increase in the latter (Figure 2D). In addition, we observed that *skn-1b* mutants are approximately 10% larger than WT (Figure 2E). This suggests that *skn-1b* mutants might ingest more *E. coli* than WT animals. To explore this further we examined food intake by quantifying uptake of fluorescently labelled OP50. If worms were fed mcherry labelled OP50 continuously (fed conditions), the guts of WT and *skn-1b* mutants contained similar amounts of bacteria (Figure 2F). However, in response to fasting and re-feeding *skn-1b* mutants accumulated more OP50 than WT, corresponding to a further increase in pumping rate under these conditions (Figures 2D and 2F). Together this suggests that *skn-1b* mutation alters feeding and quiescence associated parameters.

### Neuronal SKN-1B expression responds to specific food cues

ASI neurons detect the worm’s environment, including food cues (Bargmann, 2006). As *skn-1b* mediates food-related behaviours (Figures 1 and 2), and can contribute to DR lifespan extension (Figure 1A) (Bishop and Guarente, 2007) we examined SKN-1B expression levels in response to dietary changes. Laboratory *C. elegans* are fed a homogeneous diet of *E. coli* OP50, but can thrive on other bacterial lawns (Clark and Hodgkin, 2014). To test whether SKN-1B levels also respond to changes in food type we measured SKN-1B::GFP levels in the ASIs in *C. elegans* grown on different bacterial strains compared to OP50. SKN-1B::GFP levels were not altered when worms were cultured on *E. coli* HT115 or HB101, but increased in response to *Bacillus subtilis*, or *P. aeruginosa* (Figure 3A and 3B, Figure S7A). This induction of expression was rapid, e.g. occurring after 16hrs on *B. subtilis* (Figure 3B) and suggests that neuronal SKN-1B::GFP expression increases specifically and rapidly in response to different bacterial diets.

**Figure 3.**
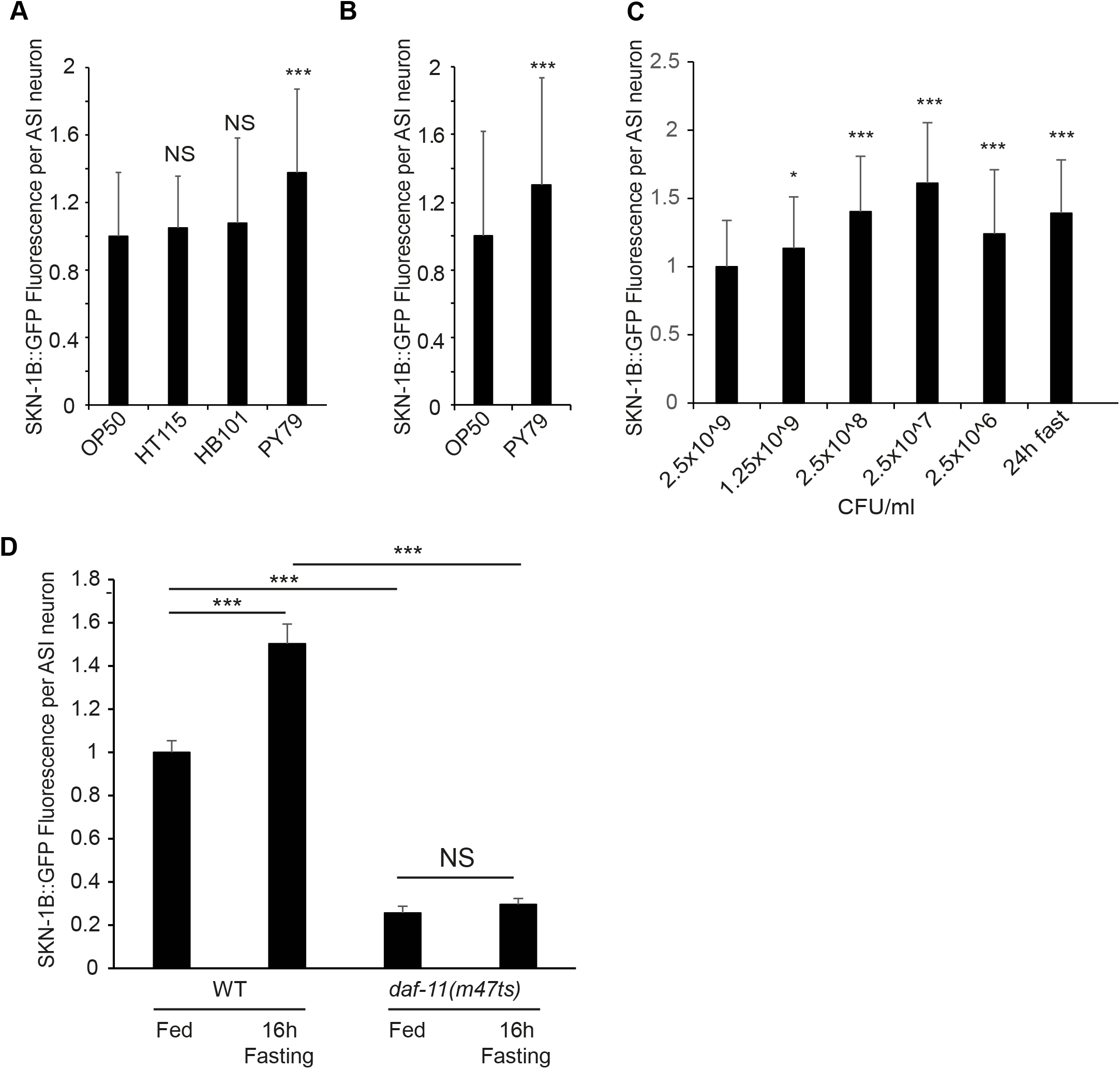
SKN-1B::GFP levels respond to nutritional cues and requires *daf-11.* **A-D)** Quantitative fluorescence microscopy of SKN-1B::GFP in response to: **A)** different bacterial strains, **B)** being switched to PY79 at the L4 stage, **C)** 16hrs bacterial dilution (Bishop and Guarente, 2007), or **C)** 16hrs fasting. For **D)** a combination of *daf-11* mutation and fasting was used. Similar results to those in **C**) had previously been observed using a SKN-1B/C::GFP transgene (Bishop and Guarente, 2007). **A-D)** Imaged at 1 day adults, each bar is the mean of 3 biological replicates ± st. dev. Two-tailed *t*-test \**p*<0.05, \*\*\**p*<0.0001, NS not significant.

As *skn-1b* contributes to DR longevity we also examined the effect of DR on SKN-1B::GFP levels. We found that diluting bacteria in liquid culture increased ASI expression of SKN-1B::GFP (Figure 3C) and that a similar increase was also observed when worms were fasted for 24hrs (Figure 3C). However, the alternative bacterial dilution DR protocol (Moroz et al., 2014) nor the *eat-2* mutation had any effect on SKN-1B::GFP levels (Figure S7B and S7C). These data suggest that neuronal SKN-1B levels respond selectively to the amount of food available.

As the behavioural effects of *skn-1b* and *daf-11* showed an interaction, we examined their relationship in respect to SKN-1B levels. Interestingly, without functional *daf-11*, SKN-1B::GFP levels were both significantly reduced, and could no longer increase in response to a 24hr fast (Figure 3D). Thus, SKN-1B requires functional *daf-11* to respond to the environment. Together with the behavioural analysis, and given the ASI expression patterns of DAF-11 (amphid opening) and SKN-1B (nucleus), this implies an epistatic relationship for these molecules, linking the external environment to SKN-1B levels and subsequent exploratory behaviour.

### SKN-1B requires TGF-β signalling to specify satiety-induced quiescence

Our data show that *skn-1b* is required in the ASIs to regulate food-related behaviours (Figures 1D-I and Figures 2A-C). One way that ASIs act is by relaying chemosensory information to the rest of the worm via secretion of neuropeptides (Bargmann, 2006). One of these, DAF-7, is the ligand of the canonical TGF-β signalling pathway, but its upstream regulators are not known (Patterson and Padgett, 2000). ASIs secrete DAF-7 under environmental conditions favourable for growth and reproduction, and DAF-7 expression is highest when worms show the high levels of quiescence (Patterson and Padgett, 2000; Wang, 2003). In addition, expression of *daf-7* in ASI has been shown to promote quiescence, whilst *daf-7* mutants do not undergo satiety quiescence (Gallagher et al., 2013; You et al., 2008). As *skn-1b* mutants’ exhibit enhanced quiescence we reasoned that *daf-7* might be a contributing factor. To test this, we generated *daf-7; skn-1b* mutants and measured their ability to undergo quiescence in response to fasting and re-feeding. In agreement with published work, WT animals showed increased quiescence following re-feeding, but *daf-7* mutants did not (Figure 4A). As before, *skn-1b* mutants spent longer than WT in quiescence (Figures 2C and 4A), but this effect proved to be completely *daf-7* dependent (Figure 4A and Figure S6B). In parallel we examined the expression of a *Pdaf-7::Venus* reporter in WT and *skn-1b* mutants. Similarly to *skn-1b, daf-7* is only expressed in ASI neurons but its expression increases in response to fasting and remains high for at least 6hrs, presumably supporting entrance into quiescence (Figure 4B and 4C). However, *skn-1b* mutants showed strongly elevated *Pdaf-7::Venus* expression in fed conditions, which barely altered in response to fasting or re-feeding (Figures 4B and 4C). Taken together, these data imply that SKN-1B inhibits satiety quiescence in response to fasting and re-feeding by suppressing *daf-7* expression and subsequently TGF-β signalling.

**Figure 4.**
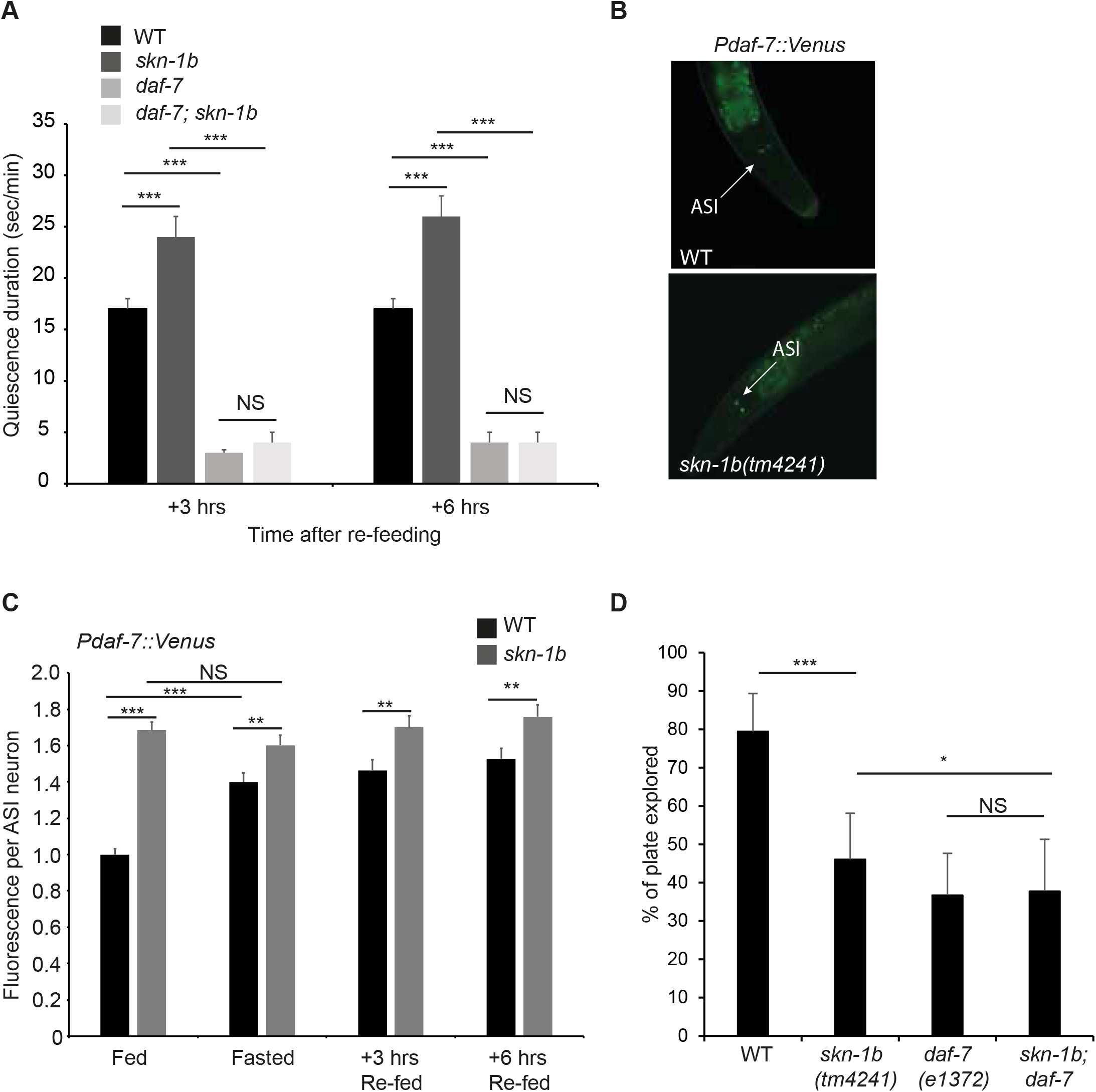
SKN-1B modulates TGF-β signalling and controls satiety. **A)** Time spent in quiescence after fasting and re-feeding. Each bar represents a mean of 3 biological replicates, ± SEM, n>9 worms per group. **B and C)** Fluorescence expression pattern **(B)**, and levels **(C)**, of *Pdaf-7::GFP* in ASIs responds to *skn-1b* mutation and food cues. In **(C)** each bar represents a mean of 3 biological replicates ± st. dev., n>230 worms per group. NS difference was found between WT samples in fasted *vs* re-fed conditions and NS difference was found between *skn-1b* samples at any point. This regulation of *daf-7* is unlikely to be direct as there is no SKN-1 binding site within 3Kb of its transcriptional start site. **D)** Quantification of exploration. Each bar is a mean of 5 biological replicates, n>44 worms per group ± st. dev. All trials shown in Figure S8A-F. For **A, C** and **D**: Two-tailed *t*-test \**p*<0.05, \*\**p*< 0.001, \*\*\**p*<0.0001, NS not significant.

*daf-7* mutants explore less than WT in fed conditions, and in this respect resemble *skn-1b* mutants (Figure 4D) (Ben Arous et al., 2009; Gallagher et al., 2013). To further investigate the behavioural epistasis relationship between *daf-7* and *skn-1b*, we examined the exploration of *daf-7*; *skn-1b* double mutants, but found that *daf-7* and *skn-1b* effects were non-additive (Figures 4D and Figure S8).

### SKN-1B modulates IIS to alter food-responsive behaviour

ASI neurons express and secrete ~40 insulin-like peptides (ILPs), at least some of which bind to the DAF-2 insulin/IGF-1-like receptor in multiple tissues (Pierce et al., 2001). rIIS leads to the de-phosphorylation and nuclear localisation of its downstream target the FOXO transcription factor DAF-16 (Antebi, 2007; Kenyon, 2010). Activation of DAF-16 has been implicated in a variety of phenotypes including behaviour, longevity, immunity and others – many of which are mediated by DAF-16 activity in the gut (Ben Arous et al., 2009; Chávez et al., 2007; Libina et al., 2003; McCloskey et al., 2017; Skora et al., 2018). To test the impact of *skn-1b* on this pathway, we examined the cellular localisation of a gut-specific DAF-16a::GFP reporter in both WT and *skn-1b* mutants. In fed conditions, *skn-1b* did not affect DAF-16 nuclear localisation (Figure 5A), but fasting for 16hrs led to DAF-16a::GFP accumulation in both WT and *skn-1b* gut nuclei (Figure 5A and Figures S9A-C). Strikingly worms lacking *skn-1b* could not maintain DAF-16::GFP in their gut nuclei after re-feeding, as WT worms do, reverting to WT levels of nuclear DAF-16::GFP within 3hrs of being returned to food (Figure 5A and Figures S9A-C) (Fletcher and Kim, 2017; Shaw et al., 2007)). Thus, *skn-1b* is required to maintain DAF-16 in the nucleus in response to fasting and re-feeding.

**Figure 5.**
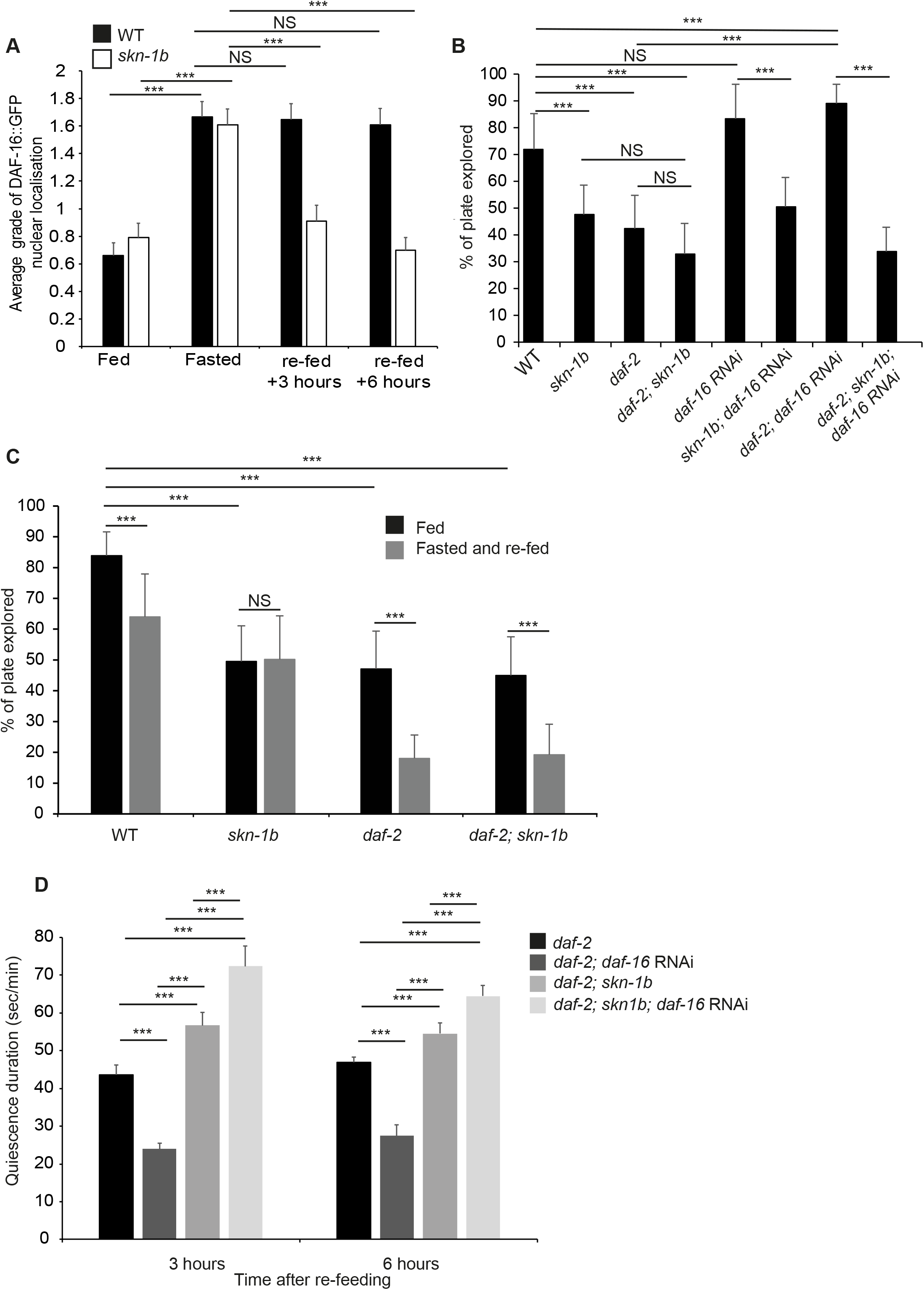
SKN-1B regulates IIS to control behaviour. **A)** Quantification of nuclear localisation, WT and *skn-1b* worms expressing *ges-1p::GFP::daf-16* (Alic et al., 2014), average grading shown. Grading system and total % of worms in each grade (Figure S9). Combined data from 3 biological replicates shown ± SEM, n>48 worms per group. **B)** Quantification of exploration. One representative of 3 biological replicates shown ± st. dev., n>10 worms per group. Similar findings were also obtained using *daf-2(e1368)* (Figure S11). **C)** Quantification of exploration in fed *vs* fasted and re-fed conditions. Worms fasted for 1hr. One representative of 3 biological replicates shown ± st. dev., n>35 worms per group. **D)** Time spent in quiescence after fasting and re-feeding. Each bar represents a mean of 3 biological replicates ± SEM, total of n>36 worms per group. For **C and D)** Similar numbers of worms from each group were observed in quiescence (Figure S7C). For **A-D**: Two-tailed *t*-test \**p*<0.05, \*\**p*< 0.001, \*\*\**p*<0.0001, NS = not significant.

Some ILPs, have *skn-1* binding sites in their promoters, making direct regulation by SKN-1B possible. One of these is *ins-7* which is expressed in several neurons, including ASIs, and the gut (Murphy et al., 2007; Oliveira et al., 2009). We observed an increase in a *Pins-7::GFP* transcriptional reporter in both the neurons and gut of *skn-1b* mutant worms (Figures S10A and S10B). INS-7 is reported to be an agonist of DAF-2 in the gut while itself being transcribed downstream of rIIS, resulting a positive feedback loop which propagates and amplifies a downregulation of IIS in this tissue (Murphy et al., 2007). Increased expression of *ins-7* in the gut might therefore explain the reduced DAF-16 nuclear localisation we observe in *skn-1b* mutants in response to fasting and re-feeding (Figure 5A).

IIS is also implicated in food-related behaviour, and *daf-2* mutants exhibit reduced exploration, similar to *skn-1b* mutants, dependent on *daf-16* (Figures 5B and S11; (Ben Arous et al., 2009; Podshivalova et al., 2017)). To try to clarify the regulatory relationship between SKN-1B and IIS we examined the relationship between *skn-1b* and *daf-16* in our behavioural assays by knocking down *daf-16* mRNA using RNAi in WT, *daf-2(e1370), skn-1b*, and double *daf-2(e1370); skn-1b* mutants. Knockdown of *daf-16* had no effect on the exploration of either WT or *skn-1b* mutants alone, but rescued the exploration deficiency of *daf-2* mutants back to WT levels (Figure 5B (Ben Arous et al., 2009; McCloskey et al., 2017; Skora et al., 2018)). Surprisingly however, *daf-16* RNAi had no effect on the exploration of *daf-2; skn-1b* mutants (Figure 5B). We also examined the relationship between *daf-2* and *skn-1b* in response to food. With food, the reduced exploration of *daf-2* and *skn-1b* mutants was non-additive suggesting that they act in the same pathway (Figure 5B). However, *skn-1b* and *daf-2* mutants respond differently to fasting and re-feeding: *skn-1b* mutant behaviour is completely unresponsive; but *daf-2* mutants respond like WT, reducing their exploration upon re-feeding, a phenotype that seems independent of *skn-1b* (Figure 2B and 5C) (Ben Arous et al., 2009; McCloskey et al., 2017; Skora et al., 2018; You et al., 2008). These data could imply either that *skn-1b* acts upstream of *daf-2* to control exploration in response to fasting and re-feeding, or that *daf-2* acts independently of *skn-1b* to control this behaviour. Overall, our data suggest that for rIIS conditions DAF-16 acts to reduce exploration and SKN-1B acts to promote it.

Our data show that *skn-1b* impacts on DAF-16 regulation in response to fasting and re-feeding, and *skn-1b* mutants cannot maintain DAF-16 in gut nuclei under these conditions (Figure 5A). rIIS increases time spent in satiety quiescence dependent on DAF-16 (McCloskey et al., 2017; Skora et al., 2018). Thus, we decided to explore whether *daf-16* contributes to the high levels of quiescence in our *skn-1b* mutants under rIIS conditions. As expected, *daf-2* mutants exhibited enhanced quiescence compared to WT (Figures 2C and 5D), but *daf-16* RNAi reduced this to WT levels, supporting the fact that *daf-16* is required for quiescence in the absence of IIS (Figure 5D) (McCloskey et al., 2017; Skora et al., 2018). *skn-1b* mutation however, increased *daf-2* quiescence, but *daf-16* RNAi did not suppress the *daf-2; skn-1b* quiescence phenotype (Figure 5D). Indeed, the quiescence of *daf-2; skn-1b; daf-16* RNAi treated animals was even higher than *daf-2; daf-16* RNAi (Figure 5D). Overall, these data suggest that SKN-1B acts to maintain nuclear DAF-16, and in doing so allows DAF-16 to promote quiescence in response to rIIS. Together with our other data, these results imply that SKN-1B acts to modulate both TGF-β and IIS in response to food, allowing the outputs of these pathways to control behaviour.

### SKN-1B controls behaviour by maintaining mitochondrial networks in muscle

Our data suggest that SKN-1B acts cell-non-autonomously to regulate behaviour. As food sensing and consumption is closely linked to physiological and metabolic homeostasis (Schmeisser et al., 2013; Sebastián et al., 2017; Weir et al., 2017), this suggests that *skn-1b* dysregulation could cause physiological and metabolic disruption to the organism. *skn-1b* is required for normal behavioural responses to fasting (Figures 1E, 1F and 2A-C), but *skn-1b* mutants are not actually starved (Figures 2D and 2E). Despite this, we noted that whilst a population of WT worms evenly distributes over a bacterial lawn, *skn-1b* mutants display a strong preference for the thicker outer edge “bordering” (Figures S12A and S12B). The edge of the lawn is considered to have reduced levels of O2 (~8%), and bordering has been associated with social behaviours, memory, temperature and starvation (Fenk and de Bono, 2017). This suggests that *skn-1b* mutants exhibit signs of starvation despite being well fed. Given that *skn-1b* mutants are unable to appropriately perceive and respond to food cues, we explored whether the physiological state of *skn-1b* mutants differs from WT.

Mitochondria are dynamic organelles that change their network morphology, balancing their fission with fusion to maximise energy production (Byrne et al., 2019; Sebastián et al., 2017; Weir et al., 2017). In worms their morphology has been shown to change in response to starvation (Hibshman et al., 2018) as well as various DR protocols (Chaudhari and Kipreos, 2017; Weir et al., 2017), and can be used to provide clues about an animal’s physiological state. In addition *skn-1* has previously been implicated in the maintenance of muscle mitochondrial networks, and anoxia-induced mitochondrial dynamics, raising the question of whether these phenomena might be mediated by *skn-1b* (Ghose et al., 2013; Palikaras et al., 2015). To explore the possibility that *skn-1b* impacts mitochondria we examined the mitochondrial networks of WT and *skn-1b* mutants expressing an outer mitochondrial membrane marker in muscle, *myo-3::GFP(mit).* We found the networks in *skn-1b* mutants to have a more fragmented appearance, covering significantly less surface area than that of the WT and the network taking on a more fragmented appearance (Figures 6A-C and Figure S12C). This is similar to the situation observed in fasted WT animals, implying that *skn-1b* mutants are, at least as far as their mitochondria are concerned, starved (Figures 6A-C). Fasting *skn-1b* mutants exacerbated these effects on the mitochondrial network, indicating that there are also other factors contributing to this mitochondrial morphology phenotype (Figures 6A-C). A similar pattern was also observed with a second mitochondrial reporter *tomm20::GFP* (Weir et al., 2017) (Figures S12D and S12E). Our data suggest that *skn-1b* contributes to maintaining muscle mitochondrial networks.

**Figure 6.**
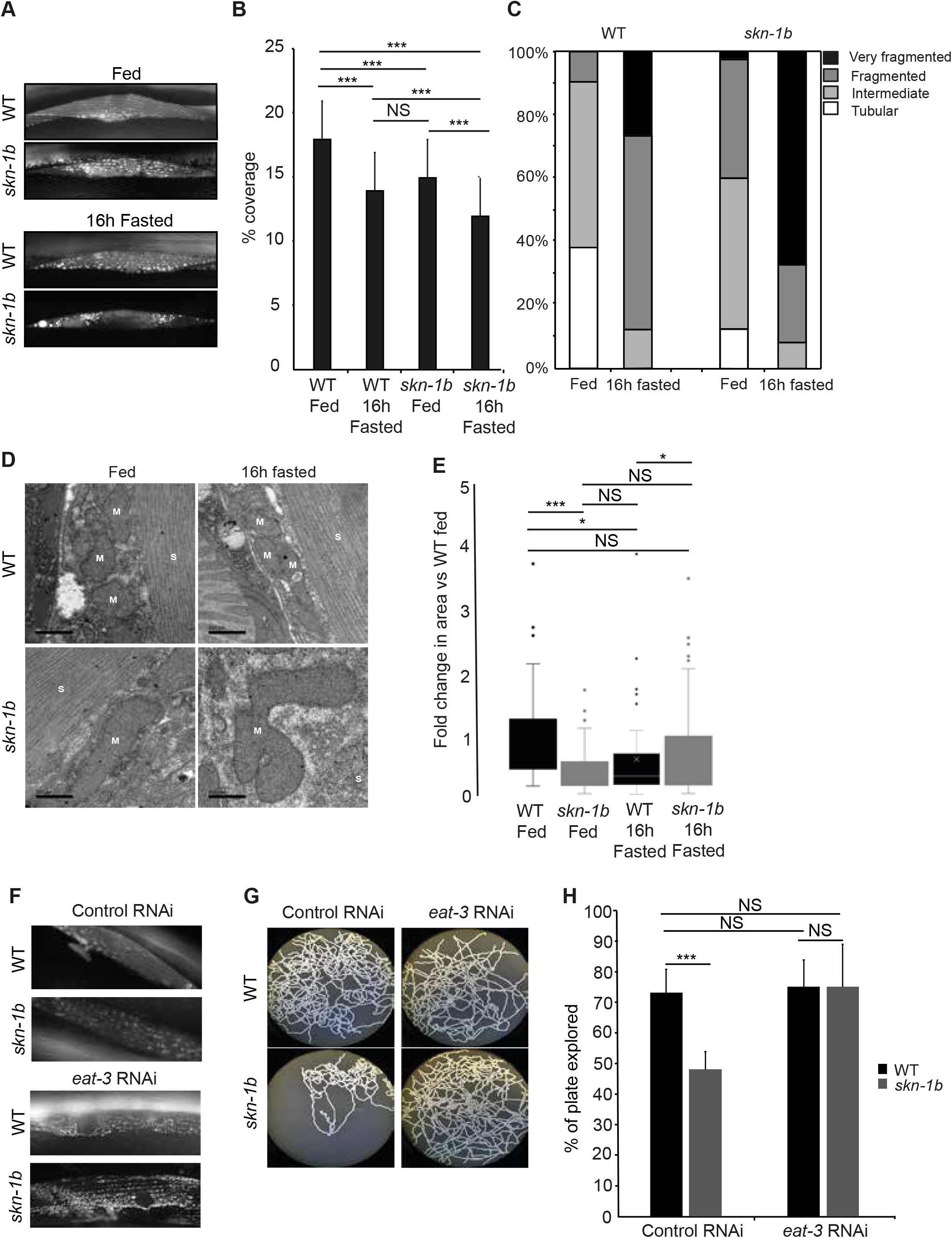
*skn-1b* contributes to mitochondrial network integrity. **A-C)** Expression and quantification of WT and *skn-1b* mutant worms expressing *myo-3::GFP(mit).* This muscle specific reporter expresses an outer mitochondrial membrane protein and hence marks all mitochondria, delineating their shape. In **B and C)** Each bar represents a mean of 3 biological replicates ± SEM, n>62 day 1 adults worms per group. The qualitative scoring system used in **C)** is shown in Figure S12E. **D)** Longitudinal sections imaged by TEM. M=mitochondria, S=sarcomere. Scale bar = 500nm. **E)** Quantification of TEM: Mitochondrial area compared to WT control. Each bar represents a mean of 2 biological replicates, n>47 images per group ± SEM. **F-H)** Effect of *eat-3* RNAi on mitochondrial networks and behaviour in WT and *skn-1b* mutants. In **F)** mitochondria are visualised using *myo-3::GFP(mit).* **G and H)** Effect of *eat-3* RNAi on behaviour in WT and *skn-1b* mutants. For all graphs: Two-tailed *t*-tests \**p*<0.05, \*\**p*< 0.001, \*\*\**p*<0.0001, NS not significant.

We then used transmission electron microscopy to examine mitochondrial morphology more closely. Muscle wall mitochondria from WT and *skn-1b* mutants were compared using sections taken from whole worms. Whist the mitochondria of fed WT animals were rounded, those in the *skn-1b* mutants, were longer and irregular, exhibiting a fused-like state (Figures 6D-E and Figure S13). This fused state was also observed in sections from WT fasted animals, supporting the idea that *skn-1b* mutant mitochondria think they are starved (Figures 6D-E and Figure S13). However, in response to fasting the mitochondrial networks of *skn-1b* mutants deteriorated further, again implying that additional factors contribute to this phenotype (Figures 6D-E and Figure S13). These data together with the fluorescence microscopy support a model whereby *skn-1b* acts to directly control mitochondrial homeostasis in response to food levels, balancing their fission and fusion.

Mitochondria require proteins in their membranes in order to fuse and fission from each other. These include *drp-1*/Drp1 that promotes fission, and *eat-3*/Opa1 and *Fzo-1/Mfn1* that promote fusion (Spurlock et al., 2020). Thus, the mitochondrial networks in *C. elegans* fed *eat-3* RNAi are more disjointed as the mitochondria are unable to fuse (Figure 6F). Mitochondrial dynamics have previously been implicated in behavioural responses (Byrne et al., 2019). So, given the behavioural role of *skn-1b* and its importance for maintaining mitochondrial networks, we tested whether the two were linked. Strikingly, we found that whilst *eat-3* RNAi had no effect on WT exploratory behaviour, it completely rescued that of *skn-1b* mutants to normal levels (Figures 6G and 6H). This supports a model whereby SKN-1B acts to regulate mitochondrial networks, which in turn control food related behaviour.

## Discussion

Ability to correctly identify a feeling of satiety impacts directly on health. For example, perception of hunger when food is plentiful, can make individuals overeat and gain excess weight, having catastrophic implications for their metabolic status and long-term health (Van Der Klaauw and Farooqi, 2015). Here we show that in *C. elegans*, the transcription factor SKN-1B, regulates satiety behaviour. SKN-1B acts in two hypothalamus-like chemosensory neurons to sense and communicate nutritional status to the rest of the organism. It then controls the animal’s behavioural responses by modulating key nutritional signalling pathways, and maintaining mitochondrial networks (Figure 7).

**Figure 7.**
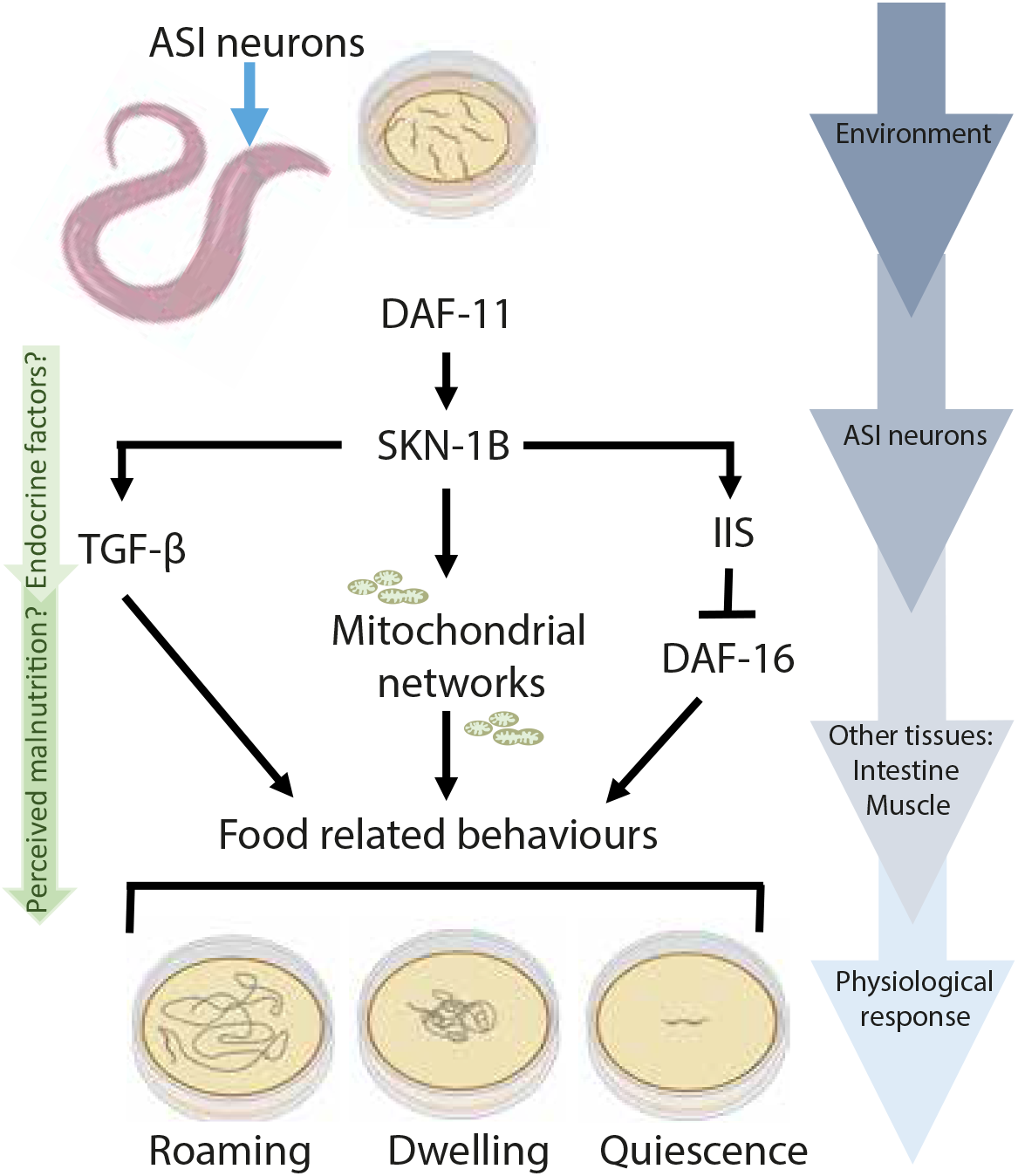
SKN-1B integrates with key nutritional signalling pathways, and regulates mitochondrial networks to control satiety-related behaviour. Food-related behaviour is controlled by interactions between food cues, SKN-1B, downstream signalling pathways (cGMP, TGF-β and IIS), and mitochondria. SKN-1B receives food cues via cGMP signalling (DAF-11). In response to fasting and re-feeding SKN-1B controls satiety quiescence: SKN-1B suppresses *daf-7* expression in the ASIs, downregulating TGF-β signalling and suppressing quiescence (Figure 4A-C). Fasting also induces DAF-16 nuclear localisation which is maintained after re-feeding to promote quiescence: SKN-1B is required for this response, possibly by acting upstream of both pathways (Figure 5A and 5D) (McCloskey et al., 2017; Skora et al., 2018). In parallel, SKN-1B is also acts to control food-related behaviour by maintaining mitochondrial networks. Overall, this study identifies neuronal SKN-1B as a novel factor in controlling satiety behaviour in response to dietary signals.

### Neuronal SKN-1/Nrf mediates the perception of food and satiety

Animals, including *C. elegans*, modulate their behaviour by integrating information about their external environment with internal cues. Our data identify SKN-1B as a novel, major regulator of food-related behaviour. SKN-1B levels respond to food availability and memories of fasting events to promote exploration in fed conditions, and suppress quiescence in response to fasting and re-feeding. We propose that SKN-1B acts as a molecular switch, allowing fine-tuning of behaviour in response to environmental change.

The constitutively nuclear expression of SKN-1B in ASI neurons (Figure 1D and Figure S3F) means that it requires the receptor type guanylate cyclase *daf-11* expressed at the amphid opening of the ASI to sense the environment. The expression pattern of *daf-11* and *skn-1b* in the ASI, the requirement of *daf-11* for SKN-1B::GFP expression, and the non-additive behavioural effects of *daf-11* and *skn-1b* strongly imply that these molecules act in the same pathway to control exploration (Figures 1H, 2A and 3D). *daf-11* has previously been mapped to act upstream of both IIS and TGF-β pathways (Hahm et al., 2009; Murakami et al., 2001; You et al., 2008), and our data identifies a new mode of *daf-11* action (Figure 7). Although *daf-11* and *skn-1b* both act to control quiescence, *daf-11* mutants exhibit decreased quiescence whilst *skn-1b* mutants have increased quiescence compared to WT (You et al., 2008) (Figure 2C). Therefore, although *daf-11* plays an important role in relation to *skn-1b’s* ties to the environment, it is likely that their behavioural responses to fasting and re-feeding are independent (Figure 7). Complete ablation of the ASIs however actually has the opposite effect to *skn-1b* mutation, reducing satiety-induced quiescence (Gallagher et al., 2013). Thus, genetic removal of SKN-1B does not “break” the neuron. Instead, we propose that specific and rapid changes in SKN-1B levels (Figures 3A-D) provide sensitivity for modulating behaviour and physiology.

### Neuronal SKN-1B modulates TGF-β and IIS to control food-related behaviour

IIS and TGF-β hormone signalling are nutritionally regulated and integral to many processes in worms and mammals. They are regulated by ILPs and NLPs, including the TGF-β ligand DAF-7. In worms they are known to control development, growth, immunity, lifespan and age-related decline (Gumienny, 2013; Lapierre and Hansen, 2012). Our data suggest that SKN-1B is a sensory switch in the ASIs, acting upstream to modulate both IIS and TGF-β signalling and allowing accurate environmental perception and behavioural control. By regulating DAF-7 in ASIs and DAF-16 in the gut SKN-1B bridges the gap between the external environment and the rest of the worm (Figures 4B, 4C, 5A and 7).

IIS is a conserved pathway for detecting food (Kimura et al., 2011) and reducing IIS using *daf-2* mutants induces quiescence dependent on DAF-16 (Gaglia and Kenyon, 2009). Without *skn-1b* however, the contribution of *daf-16* to quiescence is abolished (Figure 5D). Thus, under normal circumstances *skn-1b* allows the worm to achieve appropriate levels of quiescence for its environment (Figure 7). This interaction between *skn-1b* and *IIS/daf-16* was only revealed in the context of rIIS, and under normal conditions the two do not interact genetically to control behaviour (Figures 5B and 5D). This suggests to us that in WT *C. elegans* ILP signalling originating in the ASIs has to be “programmed” to downregulate IIS for this relationship to be important. Several ILPs could do this, but our data suggest that the insulin receptor agonist INS-7 may be important (Figures S10A and S10B). However, ILPs like INS-7 are differentially expressed in multiple tissues (Figure S10A), and have tissue specific functions making it likely that a complex intercellular network of ILP signalling will be required.

One mechanism via which DAF-16 can regulate quiescence is via food consumption. Worms carrying *daf-2* mutation eat less, and *daf-2; daf-16* double mutants consume more food (Wu et al., 2019). Our *skn-1b* mutants have reduced levels of nuclear DAF-16::GFP in their gut, which could simulate a situation comparable to *daf-16* knockdown. However, when fasted and re-fed i.e. conditions that stimulate satiety quiescence, *skn-1b* mutants exhibit higher pharyngeal pumping rates, accumulate more *E. coli* in their guts, and are slightly larger than WTs indicating that under these conditions they might be eating more (Figures 2D-F). In addition, DAF-7 levels are also higher in well fed conditions (Gallagher et al., 2013). Thus, it is possible that altered feeding parameters in *skn-1b* mutants contribute to the increase in *daf-7* reporter expression and quiescence behaviour.

### SKN-1B maintains mitochondrial networks to control food-related behaviour

We show SKN-1B acting cell non-autonomously in the gut to alter IIS, and in muscle to alter mitochondrial physiology (Figures 5A, 6A-E, Figures S12D, S12E and S13). SKN-1B supports an organised mitochondrial network, balancing fission and fusion to support energy homeostasis in both fed and fasted, re-fed conditions (Figure 6A-C, Figures S12D and S12E). Mitochondrial homeostasis is implicated in a number of processes including ageing and behaviour. A delicate balance between fission and fusion is necessary for DR to extend lifespan (Weir et al., 2017). The fused mitochondria visible in *skn-1b* mutants suggests that SKN-1B acts to control mitochondrial states (Figures 6C-E and Figure S13). The mitochondrial network observed in *skn-1b* mutants resembles that of fasted or DR worms (Weir et al., 2017), but it is unlikely that *skn-1b* mutants are physically starved (Figures 2D-F). We suggest instead, that this occurs via endocrine factors from the ASI leading to a perceived state of malnourishment, with knock-on effects for mitochondrial physiology.

Our data also shows that breaking the fused mitochondrial networks of *skn-1b* mutants using *eat-3/Opa1* RNAi is sufficient to rescue their exploratory behaviour defect (Figures 6F-H). *C. elegans* lacking the *fzo-1* or *drp-1* genes have been shown to exhibit defective movement and exploratory behaviour although *eat-3* has no impact on WT behaviour (Byrne et al., 2019) (Figures 6G and 6H). This strongly suggests that SKN-1B mediated control of mitochondrial networks is required for correct behavioural responses to food.

Our work also shows that whilst some DR protocols require *skn-1* to extend lifespan (Moroz et al., 2014), ASI specific *skn-1b* is not essential (Figures 1A and B, Tables S1 and S2) (Bishop and Guarente, 2007). Our results indicate either that the requirement for its expression varies among DR conditions or that there is redundancy for other SKN-1 isoforms in this regard. The role of SKN-1B in regulating mitochondrial networks may also influence the involvement of *skn-1b* in DR longevity (Bishop and Guarente, 2007). Mitochondrial networks are optimised for ATP production and as such generate increased levels of Reactive Oxygen Species (ROS). Fasting and DR cause mitochondrial fusion and maximise ATP production (Sebastián et al., 2017). Mitochondrial homeostasis is required for DR to extend worm lifespan (Weir et al., 2017) and, there is also evidence that small increases in ROS increase neuronal SKN-1 expression and promotes longevity (Schmeisser et al., 2013). It is possible that *skn-1b* mediated behaviours, influence the impact of DR on lifespan. Different DR protocols cause varying degrees of life extension, and *skn-1b* was required where the increases were modest. As SKN-1B subtly affects feeding this might account for these differences, potentially via changes in *skn-1b* dependent mitochondrial homeostasis.

### Potential for conservation

In mammals, linking food-status to behaviour is controlled by the neuroendocrine system, primarily the hypothalamus: Firstly by the quantity or quality of available food; and secondly by the organism’s internal state i.e. satiety signalling by gut peptides (Van Der Klaauw and Farooqi, 2015). Our data identifies SKN-1B as a key regulator of satiety quiescence, thought to mimic satiety in mammals (You et al., 2008).

Food levels also alter behaviour in fruit flies, and foraging strategies have been observed that allow adaptation to different food concentrations (Bräcker et al., 2013; Wong et al., 2009). This suggests that these processes are conserved. Nrfs have been detected in the hypothalamus (Wilson et al., 2017) and some Nrfs also have short isoforms for which functions are not known, suggesting possible conservation. Central Nervous System-specific Nrf1 knockout mice also show neuro-dysfunction phenotypes suggesting that Nrf1 plays an important role here (Kobayashi et al., 2011). Our data suggest the interesting possibility that mammalian Nrf proteins might act in the brain to regulate satiety, offering a novel pharmacological target for controlling food-related pathology.

## Acknowledgements

We thank: Queelim Ch’ng for strains and advice, Rachel McMullan for advice on food leaving assays, Simon Harvey for use of the MicroWorm tracker, David Gems for excellent support at the start of this project, and Dr Tobias Von der Haar for critical reading of the manuscript. This work was funded by a BBSRC NI award BB/R003629/1 to JMAT and NIH grants AG054215, GM122610, and DK036836 to TKB. Some strains were provided by the CGC, which is funded by NIH Office of Research Infrastructure Programs (P40 OD01044).

## Author contributions

Conceptualization, JMAT; Methodology, JMAT, NTP, MT, IGS, AFH, ZW, ME; Investigation, JMAT, NTP, MT, AFH, ZW, ME, IB, IGS; Writing Original Draft, JMAT, NTP; Writing Review & Editing, JMAT; Funding Acquisition, JMAT, TKB; Supervision, JMAT, TKB.

## Declaration of interest statement

The authors declare no competing interests.

## Methods

### Strains and cloning

Worms were cultured according as previously described (Brenner, 1974), and maintained at 20°C unless otherwise indicated. The following strains were used: N2 CGC hermaphrodite stock, GA1058 *skn-1b(tm4241)*, EU1 *skn-1*(zu67), EU31 *skn-1(zu135)*, JMT31 *daf-2(e1370)*, DR1572 *daf-2(e1368)*, DR1574 *daf-2(m1391)*, JMT32 *daf-2(e1370); skn-1b(tm4241)*, GA1060 *daf-2(e1368); skn-1b(tm4241)*, JMT5 *daf-2(e1391); skn-1b(tm4241)*, GA1017 N2 *wuEx217[Pskn-1b::skn-1b::GFP; rol-6]* (was used for all microscopy and expression analysis), GA1030 *daf-2 wuEx217*, GA1045 *daf-2; daf-16 wuEx217*, GA1034 N2 *wuEx253[Pskn-1b::GFP]*, GA1040 *daf-2 wuEx253*, GA1042 *daf-2; daf-16 wuEx253*, DA1116 *eat-2(ad1116)*, JMT7 *eat-2(ad1116); skn-1b(tm4241)*, DR47 *daf-11(m47ts)*, CB1372 *daf-7(e1372)*, JMT68 *daf-7(e1372); skn-1b(tm4241)*, JMT70 *daf-11(m47); skn-1b(tm4241)*, PR678 *tax-4(p678)*, MT1072 *egl-4(n477)*, JMT66 *skn-1b(tm4241) ukcEx15 [Pskn-1b::skn-1b::GFP; myo-3::mcherry]*, JMT67 *ukcEx16 [Pskn-1b::skn-1b::GFP; myo-3::mcherry].* JMT66 and JMT67 were used for the behavioural rescue experiment as they do not have the roller phenotype. Their expression pattern is identical to that in Figure 1D. COP1836 *knu733[wrmScarlet::skn-1b]* (created using CrispR by Knudra Biotech), GA1064 muEx227*[ges-1p::GFP::daf-16a*], SJ4103 *zIs[myo-3::GFP(mit)]*, JMT90 *skn-1b(tm4241) zIs[myo-3::GFP(mit)]*, WBM671 *wbmEx289[myo-3p::tomm20(aa1-49)::GFP::unc54 3’UTR]*, JMT76 *skn-1b(tm4241) wbmEx289[myo-3p::tomm20(aa1-49)::GFP::unc54 3’UTR]*, JMT82 *skn-1b(tm4241) muEx227[ges-1p::GFP::daf-16a];* JMT50 *drcSI7[unc-119;Pdaf-7::Venus]*, JMT75 *skn-1b(tm4241) drcSI7[unc-119;Pdaf-7::Venus]* The reporter SKN-1B::GFP reporter was made by cloning a genomic DNA fragment including 2KB directly upstream of the *skn-1b* translational start site, the *skn-1b* genomic region in front of GFP and the endogenous 3’UTR (Figure 1C). It also includes SKN-1D, but as this isoform has not been confirmed *in vivo*, we refer to it as SKN-1B::GFP. In parallel, we generated a transcriptional reporter for SKN-1B that lacked the *skn-1b* genomic region, *[Pskn-1b::GFP].* In contrast to our translational reporter, *Pskn-1b::GFP* was observed in both the ASI neurons and in the intestine (data not shown). However, as intestinal expression of SKN-1B::GFP was not observed in either of our translational reporters under any conditions tested here we conclude that SKN-1B is post translationally modified in the intestine to suppress its expression there. To examine SKN-1C specific expression we also generated a neongreen::SKN-1C CrispR strain (SUNY Biotech). *wuEx217* is used for all SKN-1B::GFP fluorescence microscopy and *ukcEx15* and *ukcEx16* were used for rescue experiments.

### Worm husbandry and Lifespan assays

Prior to experiments animals were maintained at the permissive temperature and grown for at least three generations with ample *E. coli* food source to assure full viability. Lifespan assays were performed essentially as described (Hsin and Kenyon, 1999). Survival plots and statistical comparisons (Log Rank test) were performed using OASIS2 software (Yang et al., 2011). For lifespan assays using RNAi, worms were grown on bacteria expressing the appropriate RNAi clone from the L4 stage. *E. coli* HT115 bearing the empty pL4440 vector was used as a control. NB: In some food assays worms were fed different bacterial strains. OP50 and BL21G are *E. coli* B strains, HT115, W3110 and MG1655 are *E. coli* K-12 strains, and HB101 is a B/K-12 hybrid. DA1877 is *Comamonas aquatica* and MyB11 is a *Pseudomonas sp.* encountered in the wild Bacterial isolates from (Samuel et al., 2016; F. Zhang et al., 2017; J. Zhang et al., 2017).

### Microscopy

Fluorescence microscopy: For each slide, 30-40 1 day adult worms were mounted in M9 + 0.06% tetramisole hydrochloride on a 2% agarose pad and imaged within 15 min. Imaging was conducted using a Leica DMR microscope recorded with a Leica-DFC9000GT camera. A 525/50 GFP filter was used and post-processing and quantification was performed using the Fiji distro of ImageJ. For analysing muscle fibres, ImageJ was used to apply a binary threshold to individual muscle fibres and the percentage coverage of GFP-tagged mitochondria across whole fibres calculated as in (Weir et al., 2017).

Confocal microscopy: day 1 adults were mounted on slides in CyGel (Biostatus) spiked with 0.6% tetramisole hydrochloride to immobilise. Imaging was performed using a Zeiss LSM880/Elyra/Axio Observer.Z1 confocal microscope with the airyscan acquisition mode with the 60x lens. Images were processed with ZenBlue software.

Electron microscopy: 100 L4 worms were picked into M9 buffer. M9 was then aspirated off and replaced by ~2mL 2.5% glutaraldehyde fixative in 100mM sodium cacodylate (CAB) buffer (pH7.2). Worm heads and tails were removed with a scalpel, and the bodies left overnight in fixative at 4°C. Worms were washed twice with CAB and suspended in 2% low melting-point agarose in CAB. Worms were identified in agarose suspension by dissecting scope, excised and transferred to 7mL glass vials, where they were post-fixed in 1% osmium tetroxide in CAB for 1hr at room temperature. These were washed twice in Milli-Q (10 mins each wash), and dehydrated in an ethanol series (50%, 70%, 90% for 10 mins each) followed by 100% dry ethanol (3 times, 10 mins each). Finally samples were washed 2 times (10 mins each) in propylene oxide. Agar scientific low viscosity (LV) resin was prepared fresh and mixed 1:1 with propylene oxide and added to the samples (30 mins RT). Samples were then incubated in fresh LV resin 2 times (2hrs each), embedded in LV resin by polymerising at 60°C for 24hrs. Polymerised samples were identified under a dissecting scope and individual worms were cut out and orientated on a resin block for optimal sectioning. 70nm sections were cut on a Leica EM UC7 ultramicrotome, using a Diatome diamond knife and collected onto 400-mesh copper grids (Agar Scientific). Sections were counterstained with 4.5% uranyl acetate (45 mins) and Reynolds lead citrate (7 mins). Sections were imaged on a Jeol 1230 transmission electron microscope operated at an accelerating voltage of 80kV; images acquired using a Gatan One View 4×4K digital camera.

### Behavioural assays

*C. elegans* are genetically tractable, with a characterised nervous system making them an excellent tool to study behaviour. To measure exploration, assays were performed as described (Flavell et al., 2013) (Figure S4A). 35 mm NGM plates were uniformly seeded with 200μL of saturated OP50 culture and allowed to dry overnight at room temperature. Worms were grown in uncrowded conditions to the L4 stage at permissive temperature. Individual L4 animals were placed in the centre of assay plates and transferred to 25 °C. After 16 hrs, the animals were tested to see if they were alive by gently touching them, and plates were photographed. Plates were superimposed on a grid of 3.5 mm squares and the number of squares entered by worm tracks counted. Tracks could enter a maximum of 109 squares. At least 15 (one day adult) animals per genotype were tested on three separate days using different offspring generation. Each experiment compared controls and mutants in parallel.

Food/pathogen avoidance assays were performed as described (McMullan et al., 2012). NGM plates were seeded with 100μL of bacteria culture in the centre of the plate and allowed to dry overnight. Only plates with an evenly and defined circular bacteria lawn were used for the assays. 3 well-developed adult worms from uncrowded plates were transferred to each plate. Animals were allowed to lay eggs for 4hrs at 25°C before being removed from the plates. When animals reached the L4/one day adult stage (48hrs at 25°C) plates were photographed and the numbers of worms on and off the lawn counted (Figure S6A). To measure bordering activity, these images were further analysed to stablish the % of animals on the thicker (outer ~0.5cm) part of the lawn.

Some of these assays required fasting. This was performed as described (Gallagher et al., 2013; You et al., 2008) with some modifications. Briefly, animals were maintained either on HB101 or OP50 bacteria at 20°C in non-crowed, non-starved conditions. L4 stage animals were selected and transferred either on HB101 or OP50 seeded plates for 9-12hrs until they have reached young adulthood. Then the animals were transferred with a platinum pick to 60mm NGM plates without food for 16hrs. After 16 hr of fasting, animals were transferred either on HB101 or OP50 bacteria for re-feeding. To measure satiety quiescence animals were fasted for 16hrs and then individuals were transferred to 35mm NGM plates seeded with HB101. Worms were allowed to re-feed for 3 or 6hrs before measuring quiescence. Worms found to be quiescent (cessation of movement/pharyngeal pumping) the duration of this state was measured i.e. until feeding and locomotion resumed. Pumping was measured in individual animals, videoed on food for 1 minute and the pumping rate quantified as in (Gallagher et al., 2013).

### Food intake protocol

NGM plates (3.5cm) were seeded with an overnight culture of *E.coli* OP50 expressing mCherry. Plates were stored at room temperature for two days. L4 worms were selected and either maintained on OP50 or fasted for 16hrs at 20°C. Fed or fasted worms were then placed on the fluorescent OP50 for 5 minutes and allowed to feed. Worms were imaged and the fluorescence intensity within the gut quantified.

### QPCR

RNA was isolated from adult worms after transfer of the worms to an unseeded NGM plate to remove *E. coli.* 50 - 100 worms were used for each assay. RNA was extracted using Trizol (Sigma) and cDNA synthesized using SuperScript II reverse transcriptase with oligo dT (PCR Biosystems). qRT-PCR was carried out using Fast SYBR Green Master Mix (PCR Biosystems) and the 7900 HT Fast PCR system (PCR Biosystems). Normalization of transcript quantity was carried out using the geometric mean of three stably expressed reference genes Y45F10D.4, *pmp*-3, and *cdc-42* in order to control for cDNA input, as previously described (Hoogewijs et al., 2008). Primer sequences to detect *skn-1 isoforms*, by qPCR were designed by Primerdesign as follows: *skn-1b* F: aacaggtggatcaacacggc, *skn-1b* R: ttttgcattccaatgtaggc, *skn-1a* F: agtgcttctcttcggtagcc, *skn-1a* R: gaggtgtggacgatggtgaa, *skn-a/c* F: gagagaaggggcacacgacaa, *skn-1a/c* R: tcgagcattctcttcggcag. Statistical analysis was preformed using a student *t*-test.

**Supplementary Table 1.** Data of lifespan trials using DR protocol shown in Figure 1A. Trial 3 is the representative experiment shown in Figure 1A. *eat-2* mutants were long lived in 3 out of 5 trials carried out across 20 and 25°C.

| Trial | Strain | Genotype | Mean Lifespan (days) | Temp °C | Extension (%) | p value (Log-rank) vs | n dead (total) |
| --- | --- | --- | --- | --- | --- | --- | --- |
| 1 | WT |  | 12.47 | 25 |  |  | 60 |
| 1 | GA1058 | *skn-1b(tm4241)* | 13.05 | 25 | +4.6 | WT: NS | 82 |
| 1 | DA1116 | *eat-2(ad1116)* | 13.82 | 25 | +10.8 | WT: <0.05 | 84 |
| 1 | JMT7 | *eat-2; skn-1b* | 12.18 | 25 | -2.3 | WT: NS<br>DA1116: <0.0001 | 65 |
| 2 | WT |  | 11.42 | 25 |  |  | 55 |
| 2 | GA1058 | *skn-1b(tm4241)* | 12.07 | 25 | +5.7 | WT: NS | 67 |
| 2 | DA1116 | *eat-2(ad1116)* | 11.99 | 25 | +4.9 | WT: NS | 81 |
| 2 | JMT7 | *eat-2; skn-1b* | 11.67 | 25 | +2.2 | WT: NS<br>DA1116: NS | 73 |
| 3 | WT |  | 11.04 | 25 |  |  | 84 |
| 3 | GA1058 | *skn-1b(tm4241)* | 11.6 | 25 | +5.1 | WT: NS | 75 |
| 3 | DA1116 | *eat-2(ad1116)* | 12.67 | 25 | +14.8 | WT: <0.0001 | 81 |
| 3 | JMT7 | *eat-2; skn-1b* | 11.14 | 25 | +0.9 | WT: NS<br>DA1116: <0.0001 | 74 |
| 4 | WT |  | 17.72 | 20 |  |  | 91 |
| 4 | GA1058 | *skn-1b(tm4241)* | 19 | 20 | +7.22 | WT: NS | 41 |
| 4 | DA1116 | *eat-2(ad1116)* | 24.68 | 20 | +39.3 | WT: <0.0001 | 50 |
| 4 | JMT7 | *eat-2; skn-1b* | 22.51 | 20 | +27 | WT: <0.0001<br>DA1116: <0.05 | 74 |
| 5 | WT |  | 28.42 | 20 |  |  | 60 |
| 5 | GA1058 | *skn-1b(tm4241)* | 24.14 | 20 | -12 | WT: <0.05 | 48 |
| 5 | DA1116 | *eat-2(ad1116)* | 23.57 | 20 | -17.1 | WT: <0.0001 | 58 |
| 5 | JMT7 | *eat-2; skn-1b* | 24.88 | 20 | -12.5 | WT: <0.05<br>DA1116: NS | 49 |

**Supplementary Table 2.**
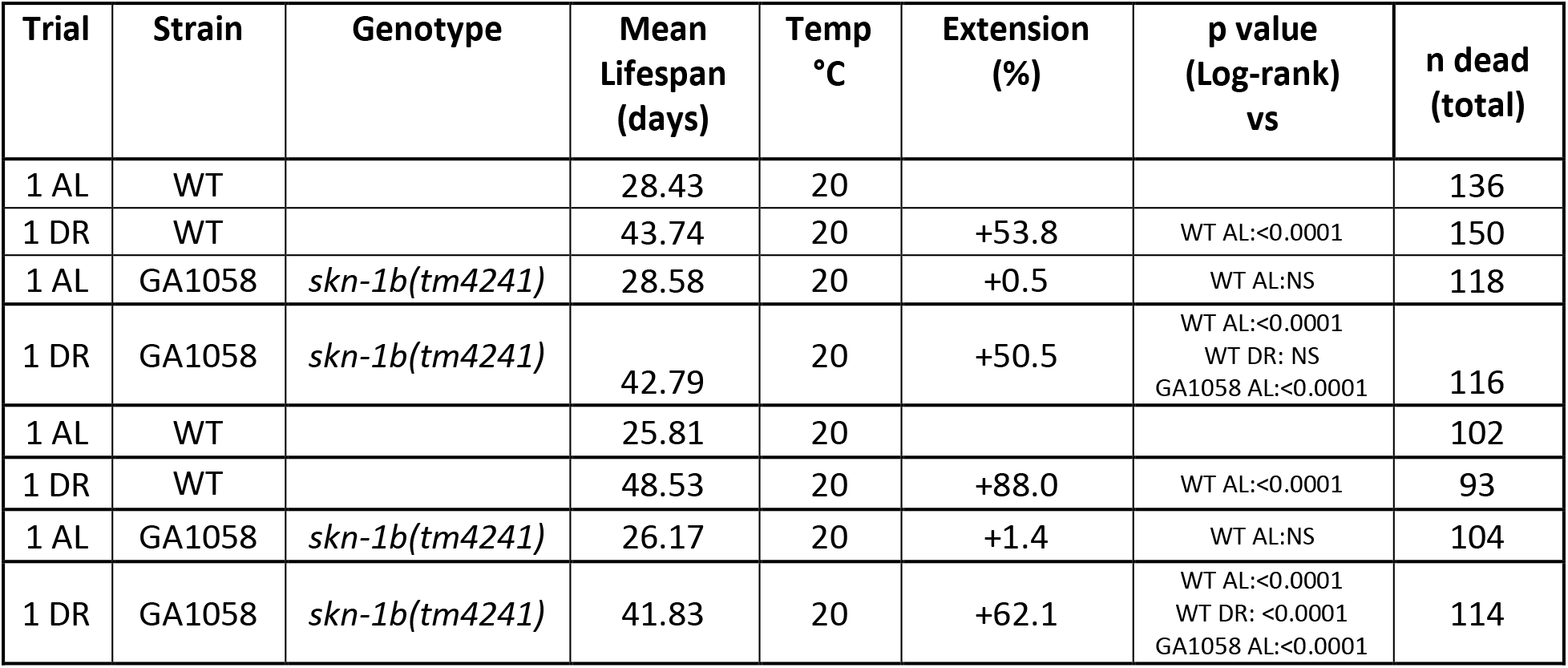
Data of lifespan trials using DR protocol shown in Figure 1B. Trial 1 is the representative experiment shown in Figure 1B.

| Trial | Strain | Genotype | Mean Lifespan (days) | Temp °C | Extension (%) | p value (Log-rank) vs | n dead (total) |
| --- | --- | --- | --- | --- | --- | --- | --- |
| 1 AL | WT |  | 28.43 | 20 |  |  | 136 |
| 1 DR | WT |  | 43.74 | 20 | +53.8 | WT AL:<0.0001 | 150 |
| 1 AL | GA1058 | *skn-1b(tm4241)* | 28.58 | 20 | +0.5 | WT AL: NS | 118 |
| 1 DR | GA1058 | *skn-1b(tm4241)* | 42.79 | 20 | +50.5 | WT AL:<0.0001<br>WT DR: NS<br>GA1058 AL:<0.0001 | 116 |
| 1 AL | WT |  | 25.81 | 20 |  |  | 102 |
| 1 DR | WT |  | 48.53 | 20 | +88.0 | WT AL:<0.0001 | 93 |
| 1 AL | GA1058 | *skn-1b(tm4241)* | 26.17 | 20 | +1.4 | WT AL: NS | 104 |
| 1 DR | GA1058 | *skn-1b(tm4241)* | 41.83 | 20 | +62.1 | WT AL:<0.0001<br>WT DR: <0.0001<br>GA1058 AL:<0.0001 | 114 |

**Supplementary Table 3.**
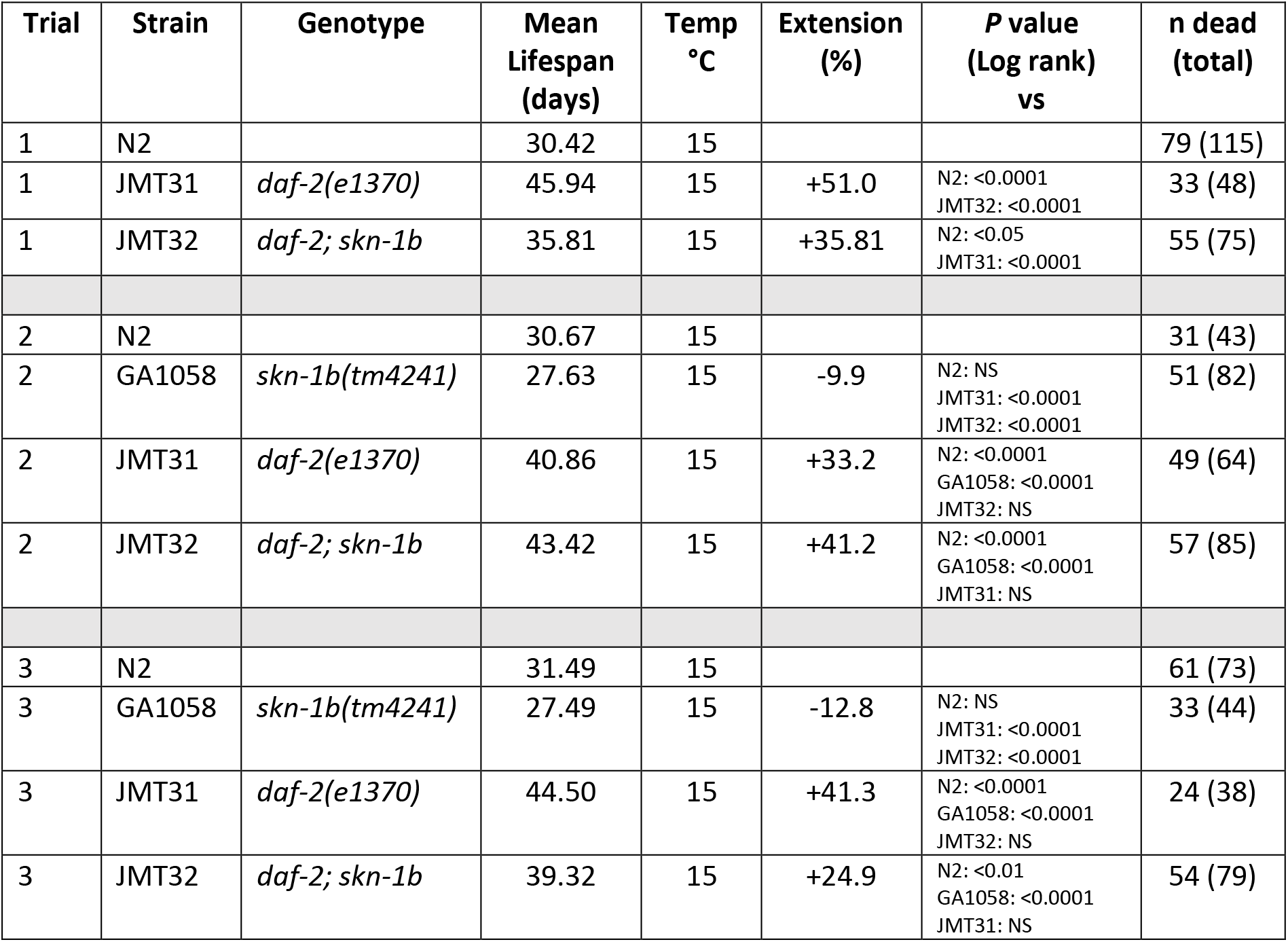
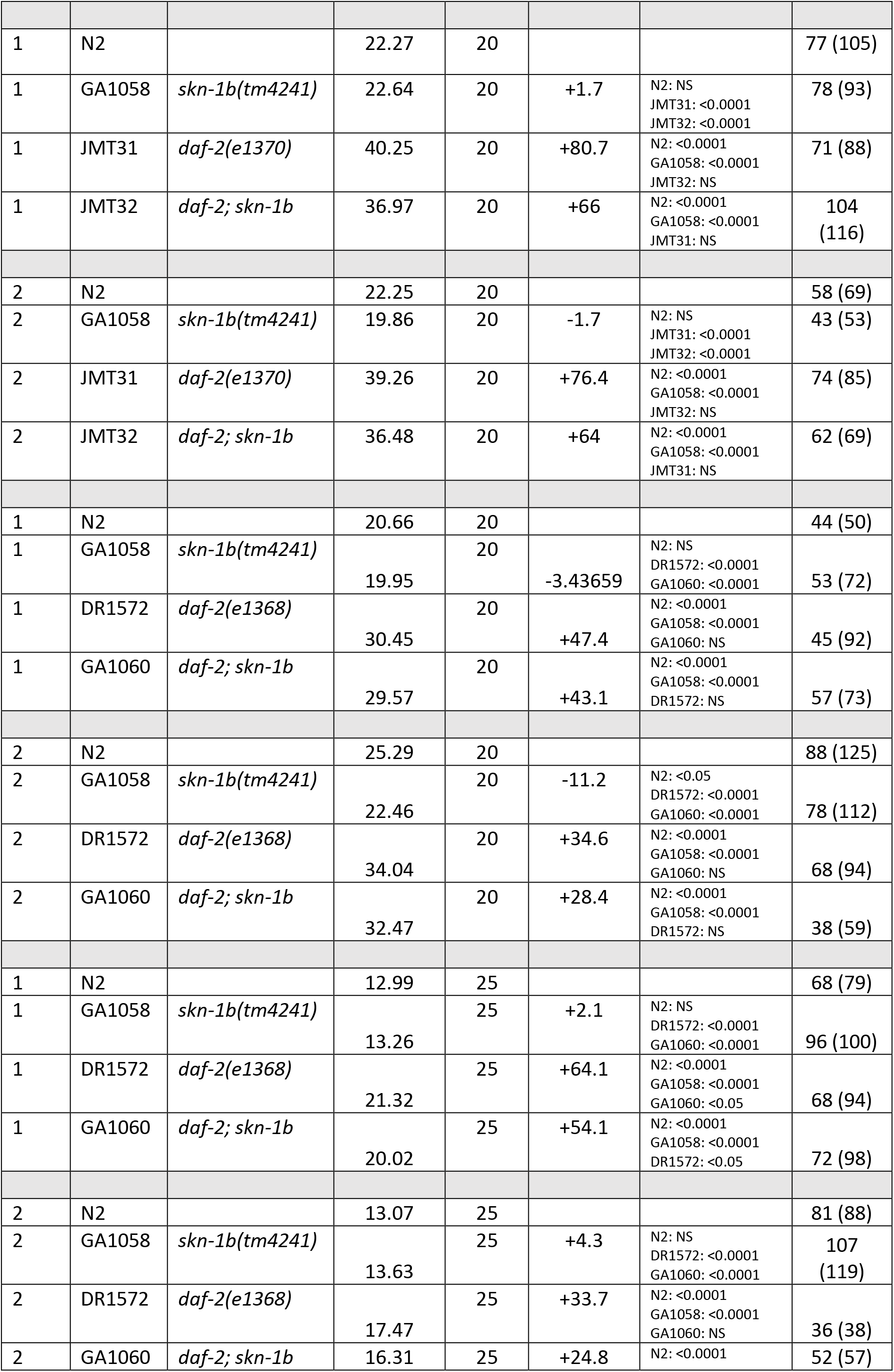
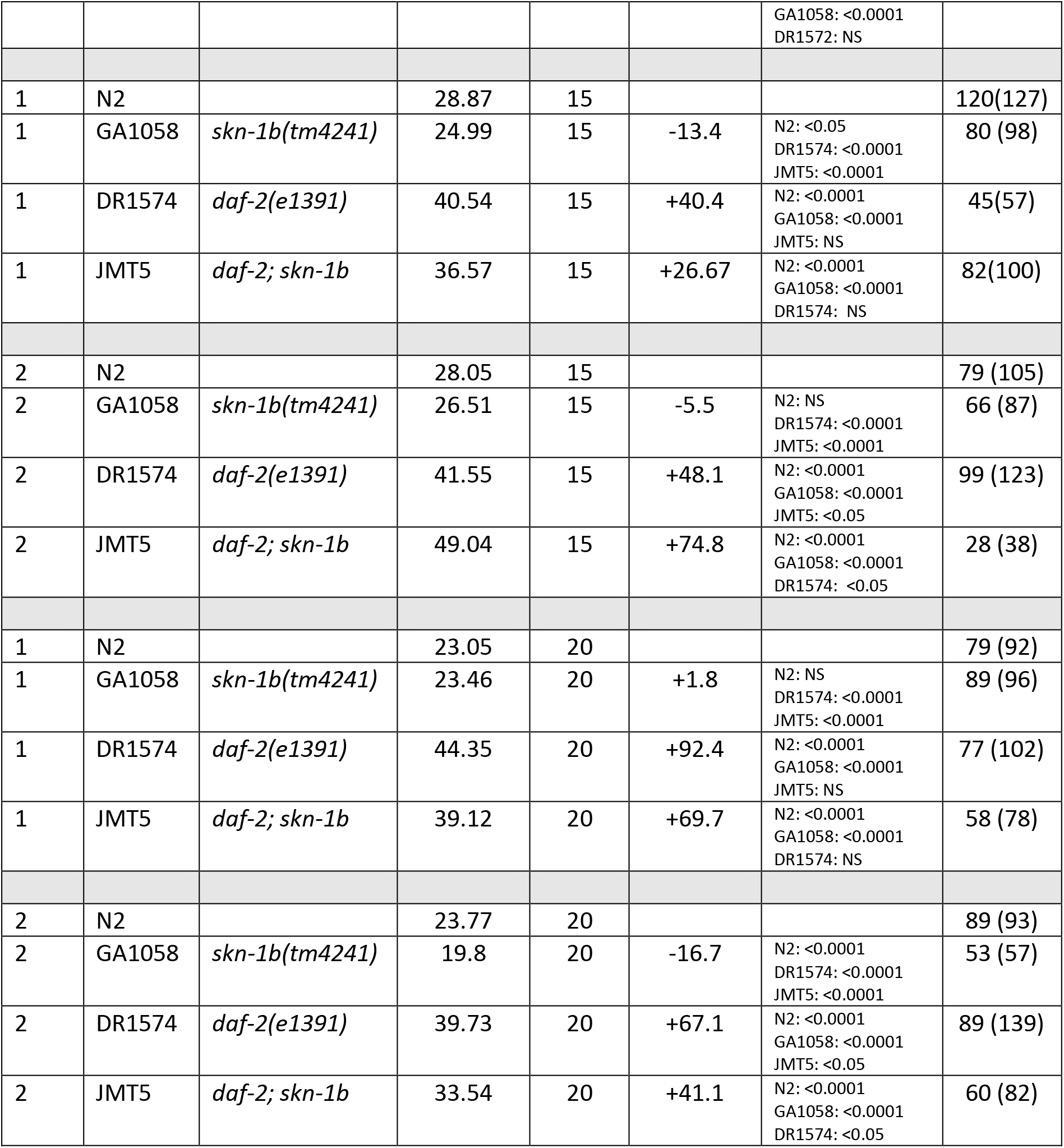
Data of lifespan trials shown in Figure S1: Figure S1A corresponds to Trial 2 in Table S1. For Figures S1B-F Trial 1 is the representative experiment in each case.

**Supplementary Table 4.** Data of lifespan trials using *daf-2* RNAi shown in Figure S2A-B. The experiment was carried out once at each temperature and the data supported the findings of the genetic experiments shown in Table S1 and Figure S1.

| Trial | Strain | Genotype | Mean Lifespan (days) | Temp °C | Extension (%) | P value (Log rank) vs | n dead (total) |
| --- | --- | --- | --- | --- | --- | --- | --- |
| 1 | N2 | Control RNAi | 28.56 | 15 |  |  | 99 (100) |
| 1 | N2 | *daf-2* RNAi | 47.74 | 15 | +59.8 | N2 C: <0.001 | 93 (100) |
| 1 | GA1058 | Control RNAi | 23.84 | 15 | -19.7 | N2 C: <0.01 | 100 (100) |
| 1 | GA1058 | *daf-2* RNAi | 47.93 | 15 | +59.3 | N2 C: <0.001<br>N2 d: NS<br>GA1058 C: <0.001 | 93 (100) |
| 1 | N2 | Control RNAi | 20.66 | 20 |  |  | 100 (100) |
| 1 | N2 | *daf-2* RNAi | 35.05 | 20 | +58.9 | N2 C: NS | 97 (100) |
| 1 | GA1058 | Control RNAi | 21.54 | 20 | +4.2 | N2 C: <0.001 | 100 (100) |
| 1 | GA1058 | *daf-2* RNAi | 35.63 | 20 | +57.9 | N2 C: <0.001<br>N2 d: NS<br>GA1058 C: <0.001 | 97 (100) |

---

**Supplementary Figure 1.**
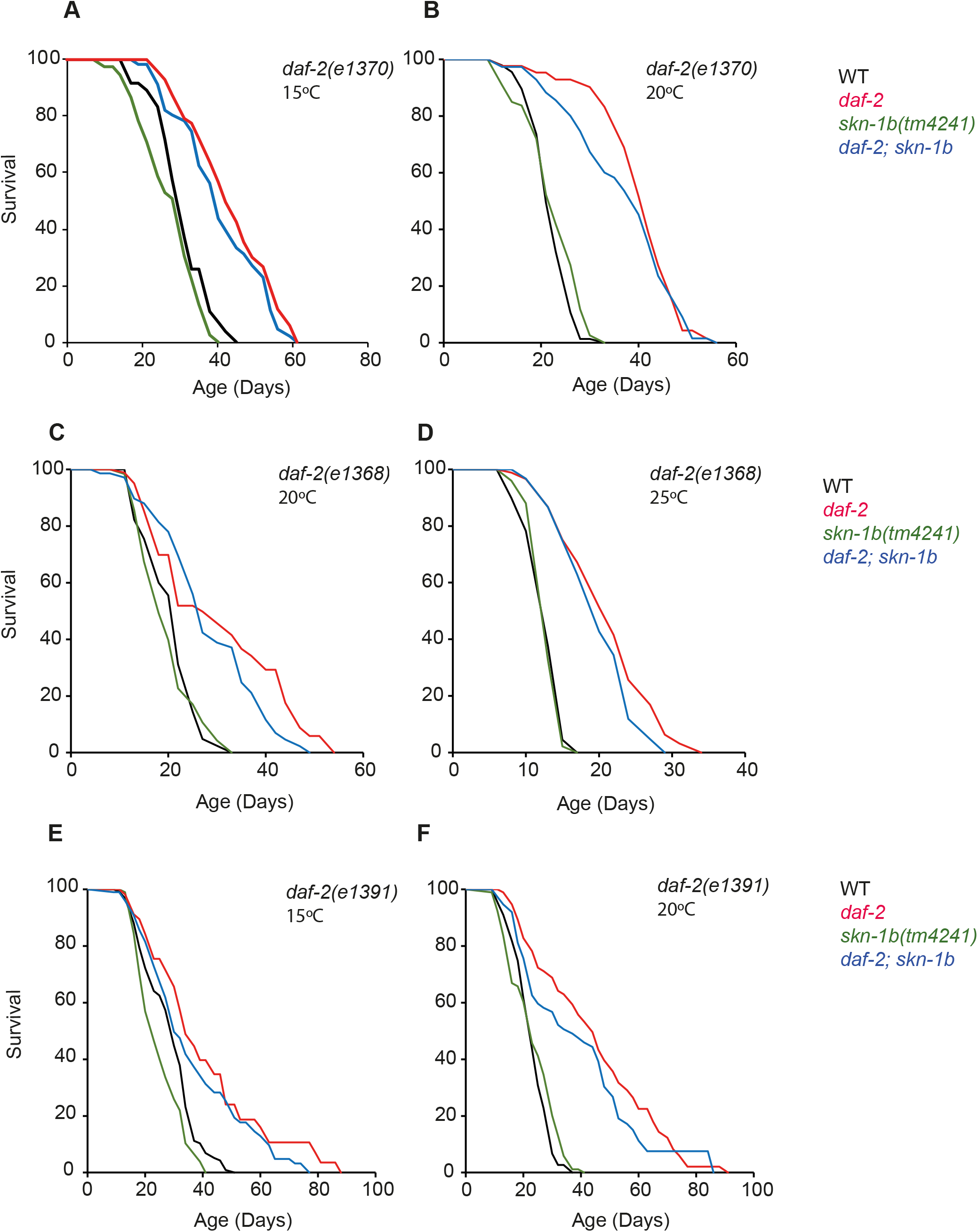
*skn-1b* is not required for WT or *daf-2* longevity. **A-F)** Survival of WT and *skn-1b* mutants in the absence and presence of *daf-2* mutation. Lower permissive temperatures were used for some *daf-2* alleles as previous work showed that *skn-1c* had a stronger suppressive effect on *daf-2* at these compared to higher temperatures (Ewald et al., 2014; Tullet et al., 2008). Representative experiments shown, individual trials are summarised with Log-Rank analysis in Table S3. NB: In a total of 13 lifespan trials we observed that *skn-1b* mutation partially suppressed *daf-2* longevity in only 4 trials (Table S3). An additional two trials using *daf-2* RNAi did not require *skn-1b* (Figure S2A-B and Table S4). We conclude that *skn-1b* does not contribute to *daf-2* longevity. In a total of 15 trials (13 on OP50, 2 on HT115) bacteria we observed a slight decrease in *skn-1b* longevity compared to WT in 4 trials (Tables S3 and S4). We conclude that *skn-1b* does not contribute to normal lifespan. *skn-1b(tm4241)* allele details (Figure S5).

**Supplementary Figure 2.**
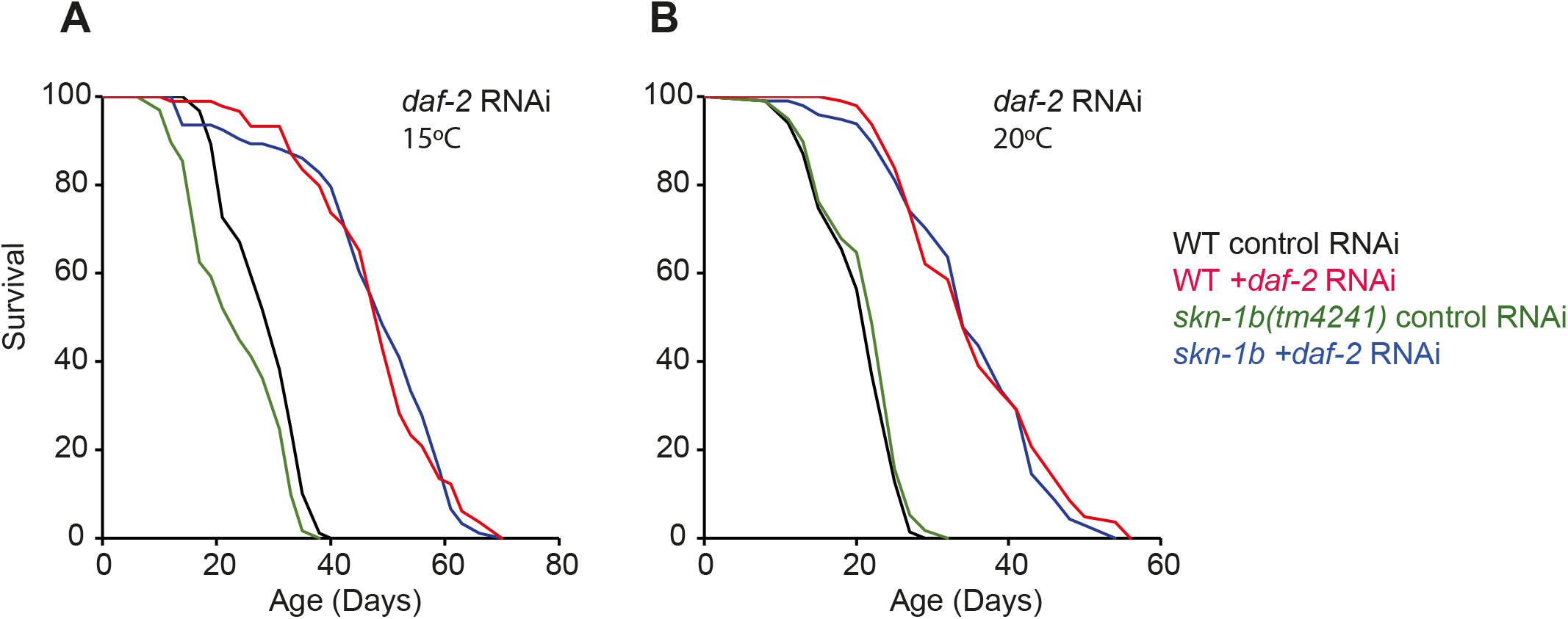
*skn-1b* is not required for *daf-2* RNAi incurred longevity. **A-B)** Survival of WT and *skn-1b* mutants in the absence and presence of *daf-2* RNAi. Full data for each trial are summarised together with Log-Rank analysis in Table S4.

**Supplementary Figure 3.**
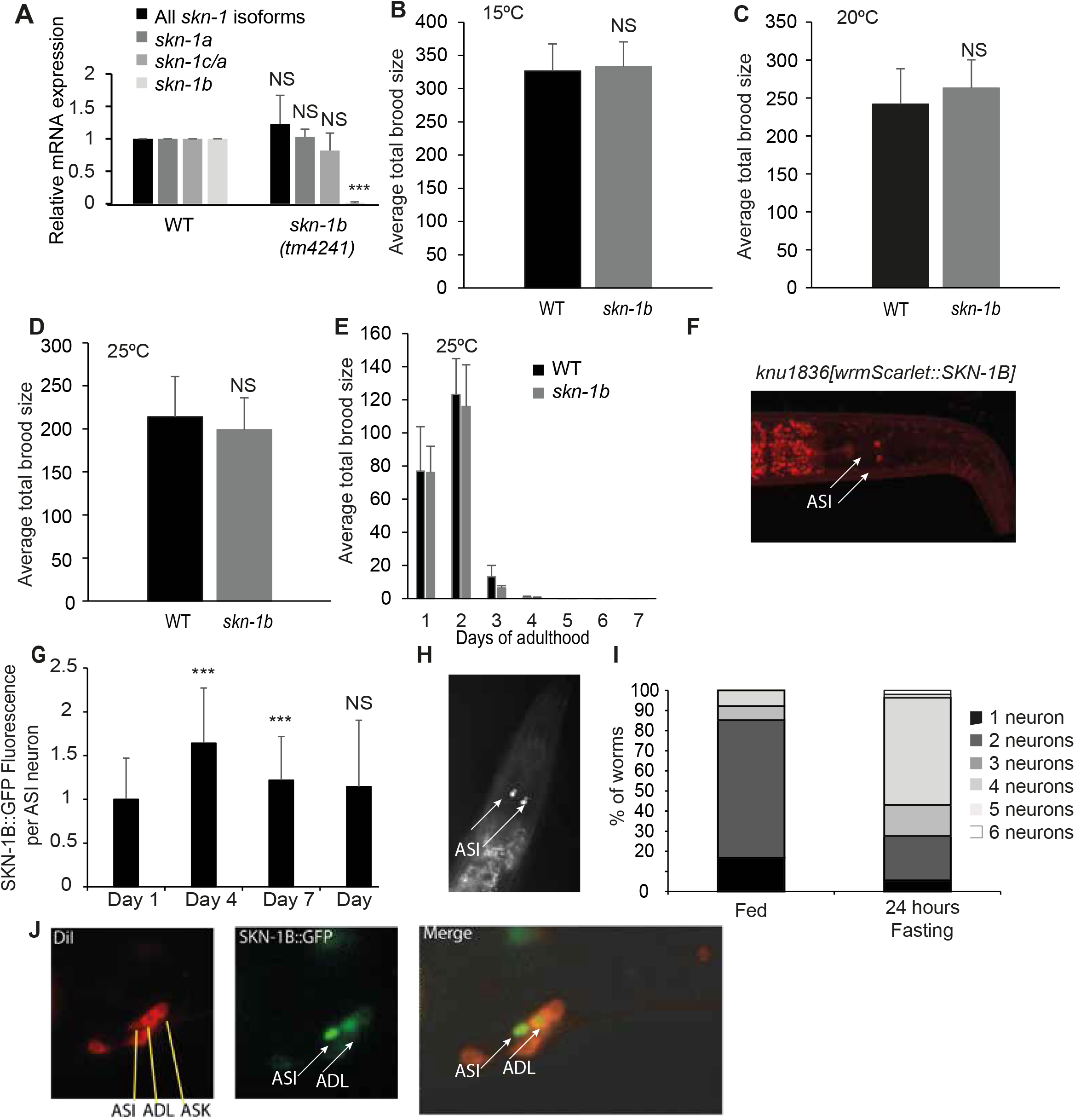
Characterisation of *skn-1b(tm4241)* and SKN-1B::GFP. **A)** Expression levels of *skn-1* isoforms in WT and *skn-1b(tm4241)* determined by Q-PCR. Combined data from 6 biological replicates shown. Error bars show st. dev. Two-tailed t-test compared to WT control *p<0.05, **p< 0.001, ***p<0.0001, NS not significant. **B-D)** Brood size of WT and *skn-1b* mutants at three different temperatures. *skn-1b* mutants are fully fertile, so can be maintained as homozygotes. Combined data from 3 biological replicates shown, n>30 worms per group. Error bars show st. dev. Two-tailed t-test *p<0.05, **p< 0.001, ***p<0.0001, NS not significant. **E)** Age-specific fecundity in WT and *skn-1b* mutants. **F)** Expression pattern of the Scarlet::SKN-1B reporter in day 1 adults under fed conditions shows SKN-1B in ASI neurons. Our lab also generated an endogenous NeonGreen::SKN-1C reporter, but cannot detect expression of SKN-1C in neurons (data not shown). No significant differences were observed using a two-tailed t-test on any day between genotypes. **H-J)** SKN-1B::GFP is expressed in additional neurons in response to bacterial deprivation. **H)** SKN-1B::GFP observed in additional neurons in response to fasting. **I)** Quantification of the number of visible neurons in SKN-1B::GFP expressing worms in response to fasting. A total of 52 fed and 64 fasted worms were examined. **J)** DiI staining confirms SKN-1B::GFP in the ASI neurons and identified two of these additional neurons (counted in **Figure S3I**) as being the ADLs. Recently, others have identified SKN-1B in AIY neurons as a regulator of chemosensory processes and behaviour, showing that animals lacking the *skn-1a, c* and *b* do not chemotax towards NaCl, butanone or temperature, or move towards thicker bacterial lawns (as a WT worms would) (Wilson et al., 2017). We tested our *skn-1b* specific mutant in a NaCl chemotaxis assay and got similar results (data not shown) but have no evidence that SKN-1B is endogenously expressed in the AIYs in fed conditions. It is possible that SKN-1B signals from the ASI - AIY neurons to mediate this response or that some SKN-1B::GFP expressing neurons in fasted conditions are AIYs. As the no-food conditions in NaCl chemotaxis assays is sufficient to induce SKN-1B expression in these cells, this could mediate the effects.

**Supplementary Figure 4.**
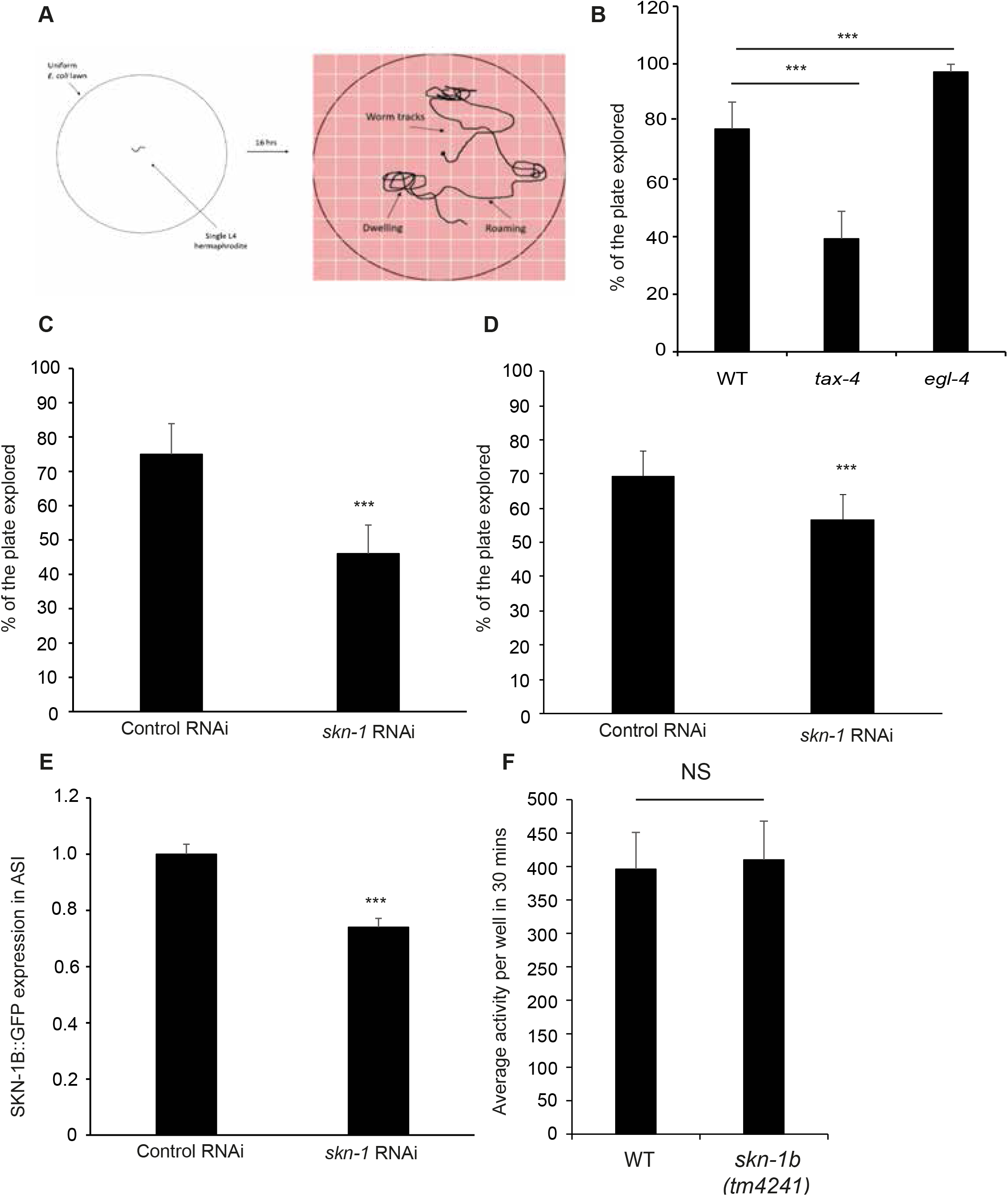
Role of *skn-1b* in regulating exploratory behaviour. **A)** Cartoon showing setup of exploration assay as in (Flavell et al., 2013). Extended dwelling or roaming compared to WT behaviour can be quantified by counting the number of squares that a worm traverses over 16hrs. Food is spread evenly and continuously on the plate. **B)** Control experiments for exploration assay. Time spent in roaming and dwelling states depends on integrating internal neuro-modulatory cues with external sensory cues. The absence of such sensory transduction leads to extended dwelling as observed in the *tax-4* mutant, *tax-4* encodes a cyclic nucleotide-gated channel subunit, in contrast, mutants with constitutive sensory input, such as the *egl-4*, which encodes a molecule with cGMP protein kinase activity, exhibit extended roaming (Fujiwara et al., 2002). Representative experiment of 3 biological replicates shown, n<15 worms per group ± st. dev. **C and D)** Quantification of exploratory behaviour in response to *skn-1* RNAi fed at either the L1 **(C)** or L4 stage **(D)**. Mean plate coverage of n>23 individual worms per group ± st. dev., one representative experiment of 3 biological replicates shown. For **B–D)** Two-tailed t-test NS non-significant. Two-tailed t-test *p<0.05, **p< 0.001, ***p<0.0001, NS not significant. **E)** Neurons are relatively resistant to RNAi (Timmons et al., 2001), but quantitative fluorescence microscopy shows that *skn-1* RNAi from the L1 stage reduces SKN-1B::GFP in ASIs by ~30%. The smaller difference between exploration in WT and *skn-1b* mutants in **(D)** likely reflects the RNAi knockdown from L4 being less complete. The *skn-1* RNAi clone used in **C-E** targets all *skn-1* isoforms (Tullet et al., 2008). Pooled data from 3 biological replicates shown, n>100 individual worms per group. Two-tailed t-test ***p<0.0001. F) *skn-1b* mutants display normal thrashing activity in liquid. Average of 3 biological replicates shown, n>33 individual worms per group.

**Supplementary Figure 5.**
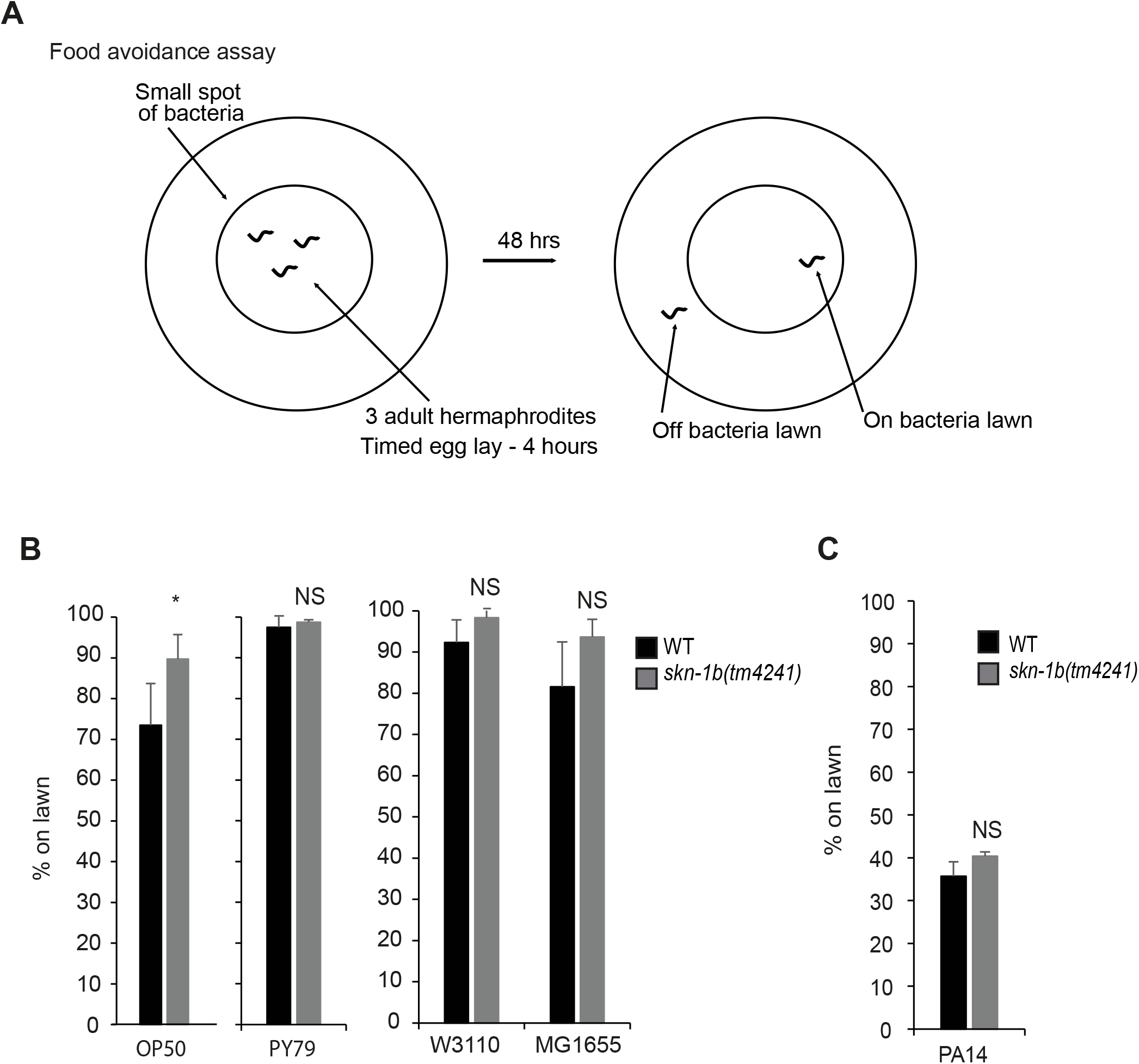
Effect of *skn-1b* mutation on food avoidance behaviour. **A)** Cartoon showing setup of food avoidance assay as in (McMullan et al., 2012). The percentage of worms on a lawn of bacteria is determined in conditions where worms have a choice whether to be on or off the lawn. **B-C)** Quantification of worms on different bacterial lawns (if given a choice to leave). Other strains tested are shown in Figure 2F. Each bar represents a mean of 3 biological replicates with ~100 worms per trial ± st. dev. Two-tailed t-test *p<0.05, **p< 0.001, ***p<0.0001, NS not significant. For **B-C)** bacteria were allowed to proliferate in each case, and no antibiotics or FUDR were present (see star methods). NB: Our assay measures satiety quiescence following fasting, as this offers an easily quantifiable behaviour. However, satiety quiescence also occurs cyclically between foraging and dwelling when worms are fully fed (although this is harder to measure without an automated tracker). During our studies we observed that while *skn-1b* mutants always preferred the bacterial lawn regardless of the food type, WT worms could be tempted to spend more time on certain bacteria (Figure 2F). This may be because *skn-1b* mutants cannot sense differences between bacterial food types; or what we are observing is an increase in satiety quiescence in fed conditions. There are examples of strains, e.g. *daf-2* mutants that quiesce more in both fed, and fasted/re-fed conditions, suggesting that the enhanced ability to enter the quiescent state is maintained across food conditions (McCloskey et al., 2017; Skora et al., 2018). In the future it will be important to ascertain this for SKN-1B.

**Supplementary Figure 6.**
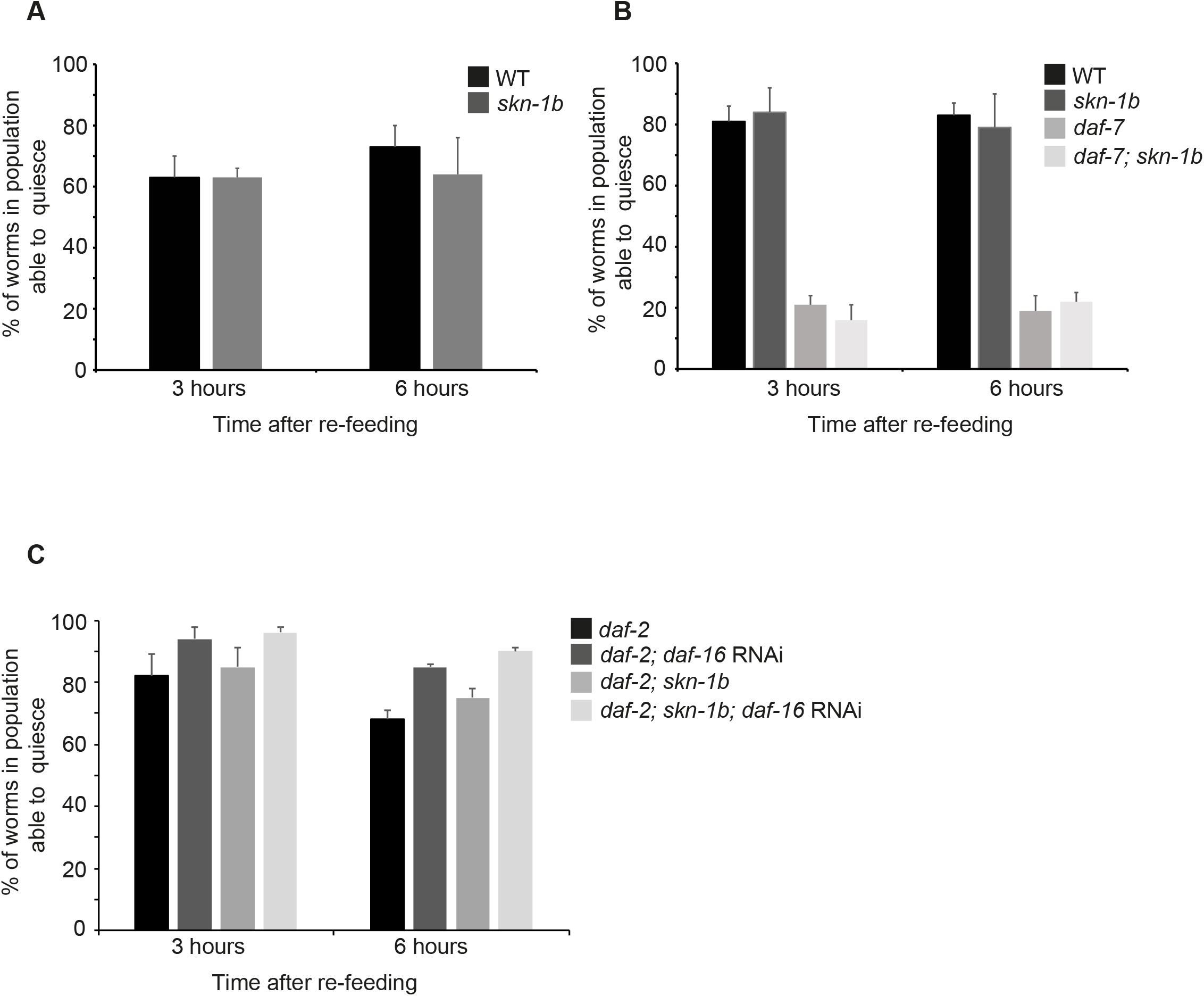
Role of *skn-1b* in mediating entry to quiescence. **A-C)** % worms spending time in quiescence 3 or 6hrs after fasting/re-feeding. Each bar represents a mean of 3 biological replicates ± SEM with n>36 worms per group. Due to the nature of the assay, satiety quiescence is not observable in every worm in an experiment, particularly in mutant strains that exhibit low levels of quiescence such as *daf-7.* Similar numbers of worms from WT and *skn-1b* mutants were observed in quiescence **(Figure S6B)** but fewer *daf-7* and *daf-7; skn-1b* mutants entered quiescence **(Figure S6B)**. Thus, the *daf-7* data in **Figure 4A** is likely to be an over-representation of the actual level of satiety quiescence within the population.

**Supplementary Figure 7.**
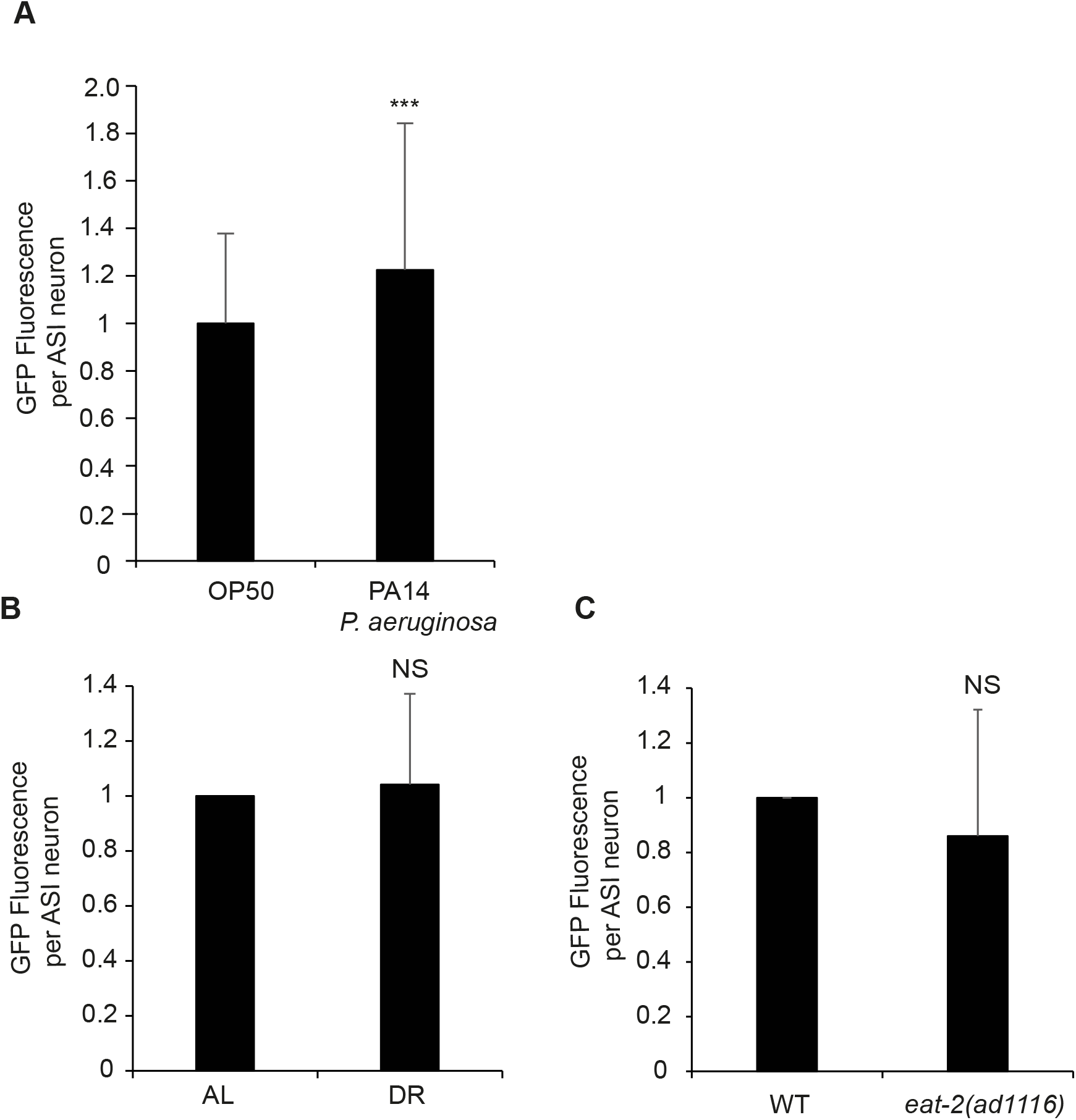
SKN-1B expression and response to environmental change. **A-C)** Quantitative fluorescence microscopy of SKN-1B::GFP expression in response to pathogenic bacteria **(A)**, an alternative DR protocol (Moroz et al 2014) **(B)**, or *eat-2* mutation **(C)**. For **(A and C)** bacteria were allowed to proliferate in each case, in **B)** antibiotics were present (see methods). For **A-C)** Error bars show st. dev. Two-tailed t-test compared to day 1 expression levels *p<0.05, **p< 0.001, ***p<0.0001, NS not significant.

**Supplementary Figure 8.**
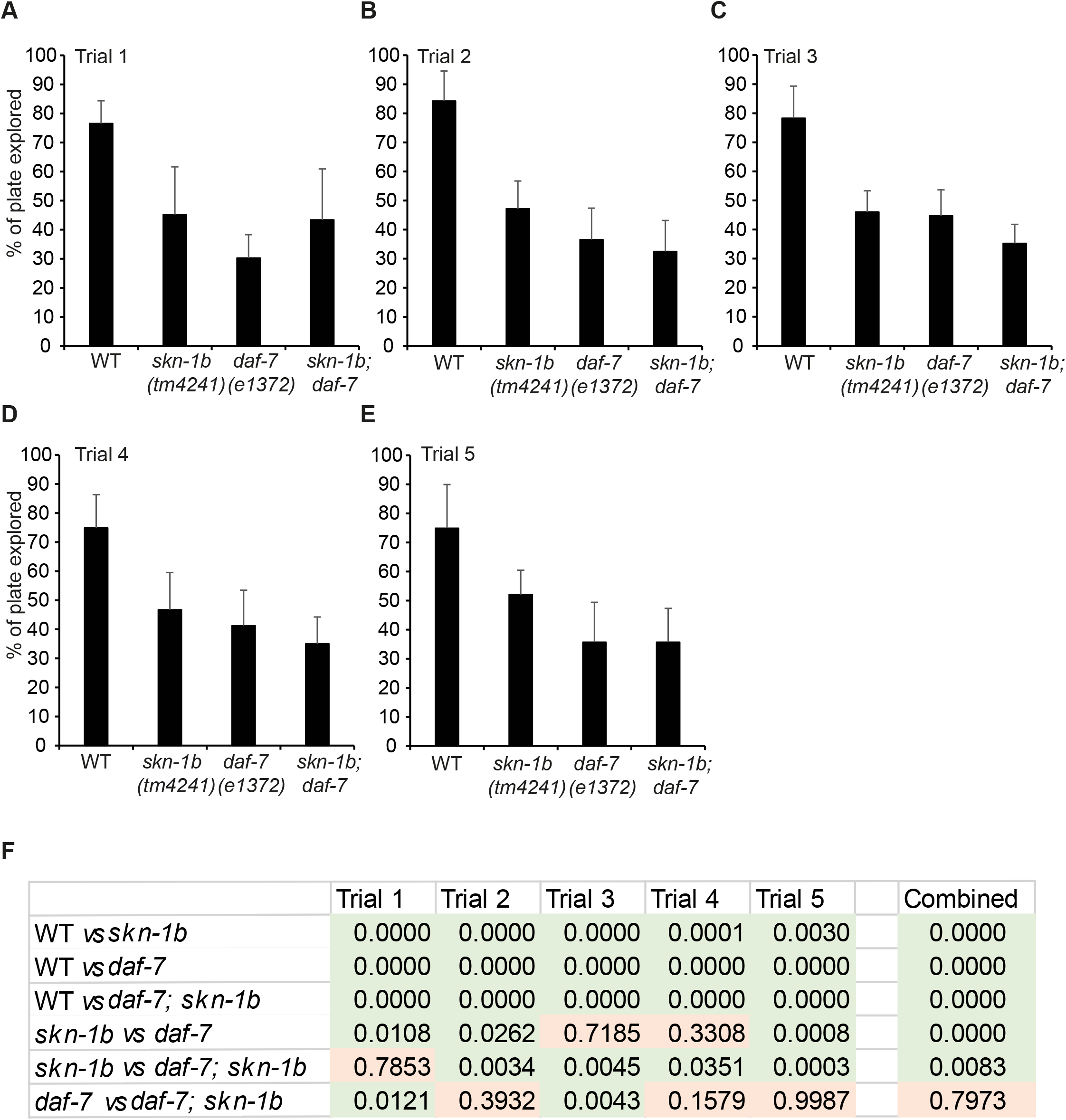
Epistasis of *daf-7* and *skn-1b* in regulating exploration. **A-E)** Individual exploration assays combined in **Figure 4D**. We reasoned that if each of the two genes regulate different behaviours independently, then the effects of *daf-7* and *skn-1b* on behaviour should be additive. However, the exploration of *daf-7* and *daf-7; skn-1b* worms was not significantly changed in 4 out of 5 trials i.e. not additive effect. In each experiment the mean plate coverage of n>8 individual worms per group is shown ± standard deviation. Two-tailed t-test *p<0.05, **p< 0.001, ***p<0.0001, NS not significant. **F)** Statistical analysis of each individual and combined trial(s). Comparisons highlighted in green are significant (two-tailed t-test p<0.05), and those in orange are NS.

**Supplementary Figure 9.**
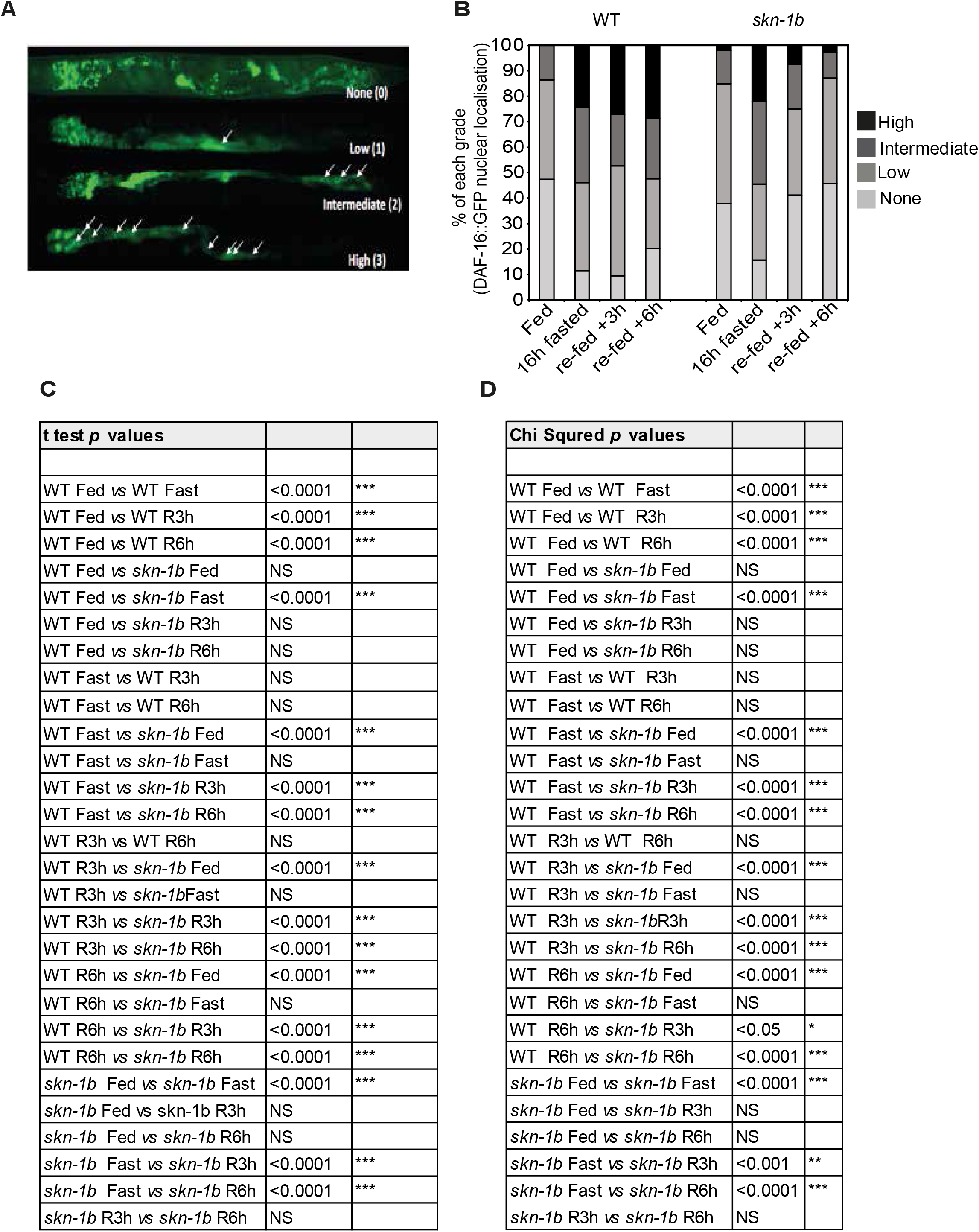
*skn-1b* alters DAF-I6::GFP nuclear localisation in response to diet changes. **A)** Scoring system for DAF-l6a::GFP nuclear localisation in the gut nuclei. Nuclear localisation was graded by a four-point system; 0=none, 1=low, 2=intermediate, 3=high. Nuclear grading was carried out by a combination of the quantity of punctate gut nuclei as well as the fluorescence intensity of these nuclei. **B)** Quantification of the grading of the DAF-16a::GFP nuclear localisation in both WT and *skn-1b* mutants under fed, fasted, fasted/re-fed for 3hrs or 6hrs. **C)** Full statistical analysis using two tailed t-test for the average grading of DAF-16a::GFP nuclear localisation shown in **Figure 5A**. **D)** Full statistical analysis determined by chi-squared test of DAF-16 nuclear localisation data shown in **Figure S9B**.

**Supplementary Figure 10.**
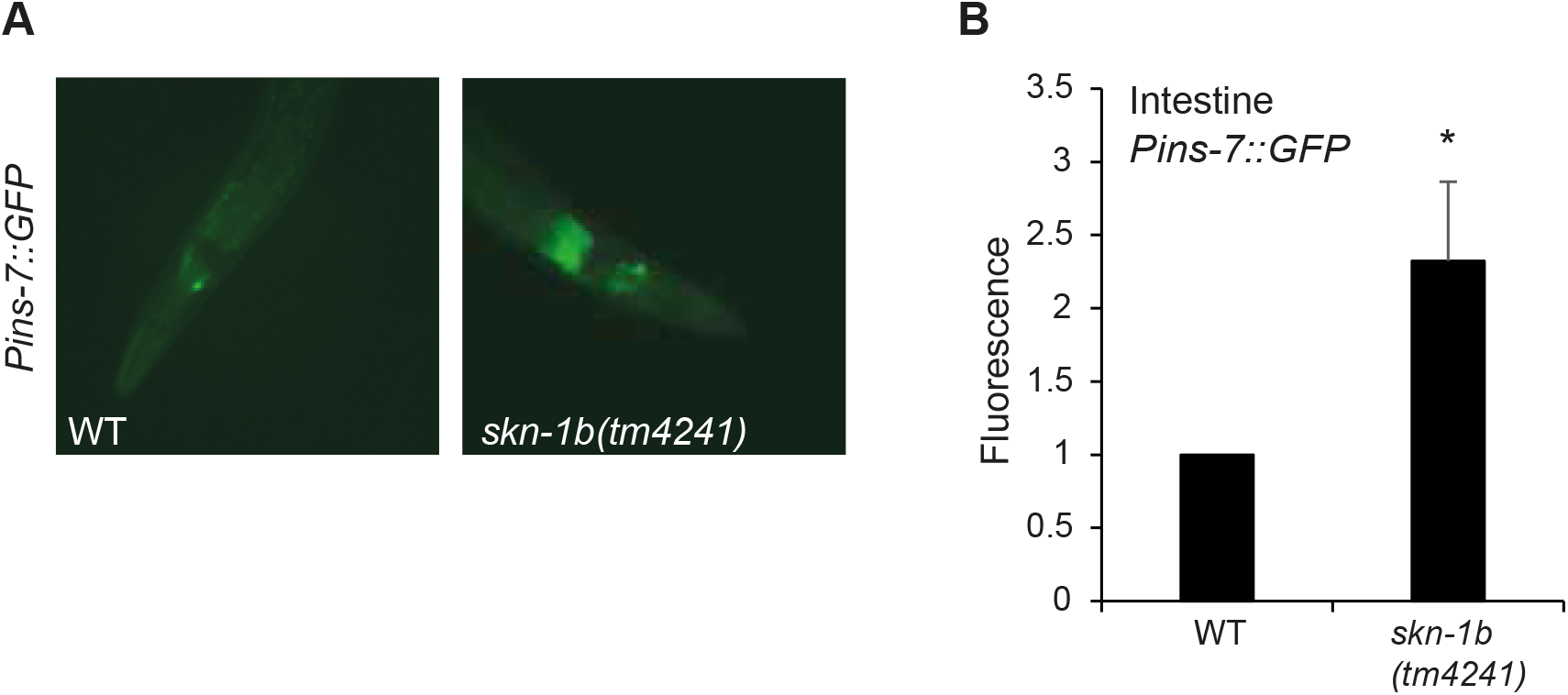
*Pins-7::GFP* levels are altered in *skn-1b* mutants. **A)** Representative images showing *Pins-7::GFP* expression in WT and *skn-1b* mutants. Expression was visible in various neurons and the gut. **B)** Quantitative fluorescence microscopy of *Pins-7::GFP* in the gut. Neuronal *Pins-7::GFP* levels were not quantified as its expression in multiple neurons made their individual identification difficult.

**Supplementary Figure 11.**
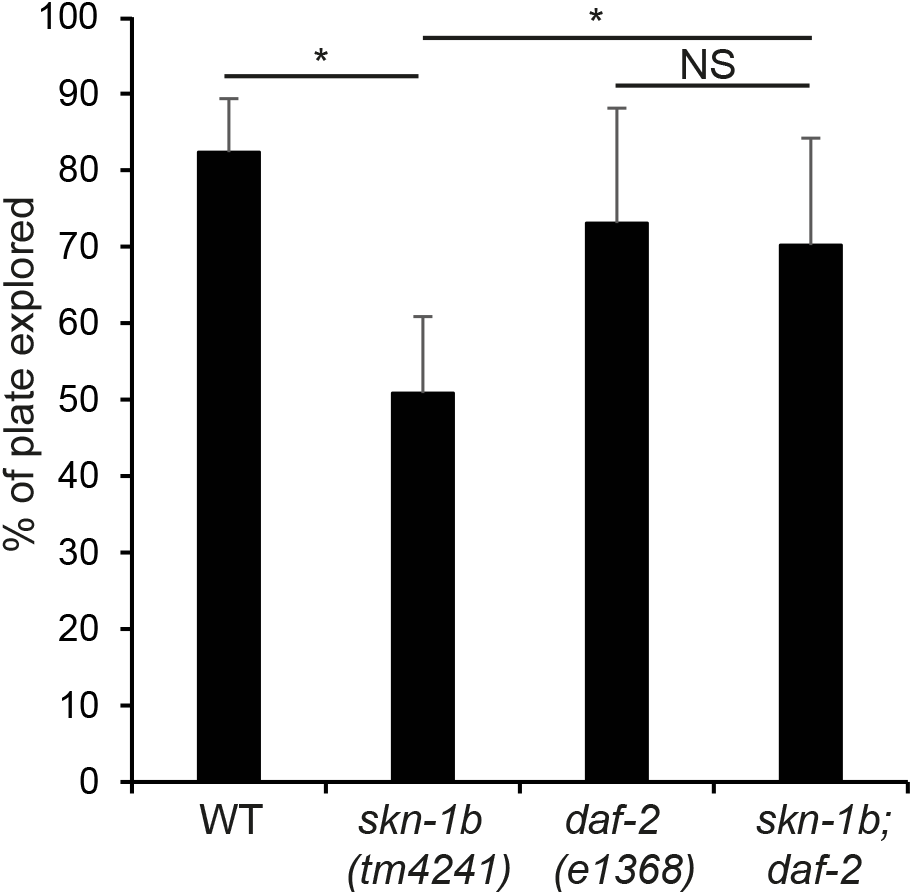
Interaction of *skn-1b* and IIS for exploratory behaviour. Quantification of exploration. One representative experiment of 3 similar biological replicates shown ± st. dev., n>10 worms per group. Two-tailed t-test *p<0.05, **p< 0.001, ***p<0.0001, NS not significant. *daf-2(el368)* caused a milder exploratory defect than *daf-2(e1370)* **(Figure 5B)**.

**Supplementary Figure 12.**
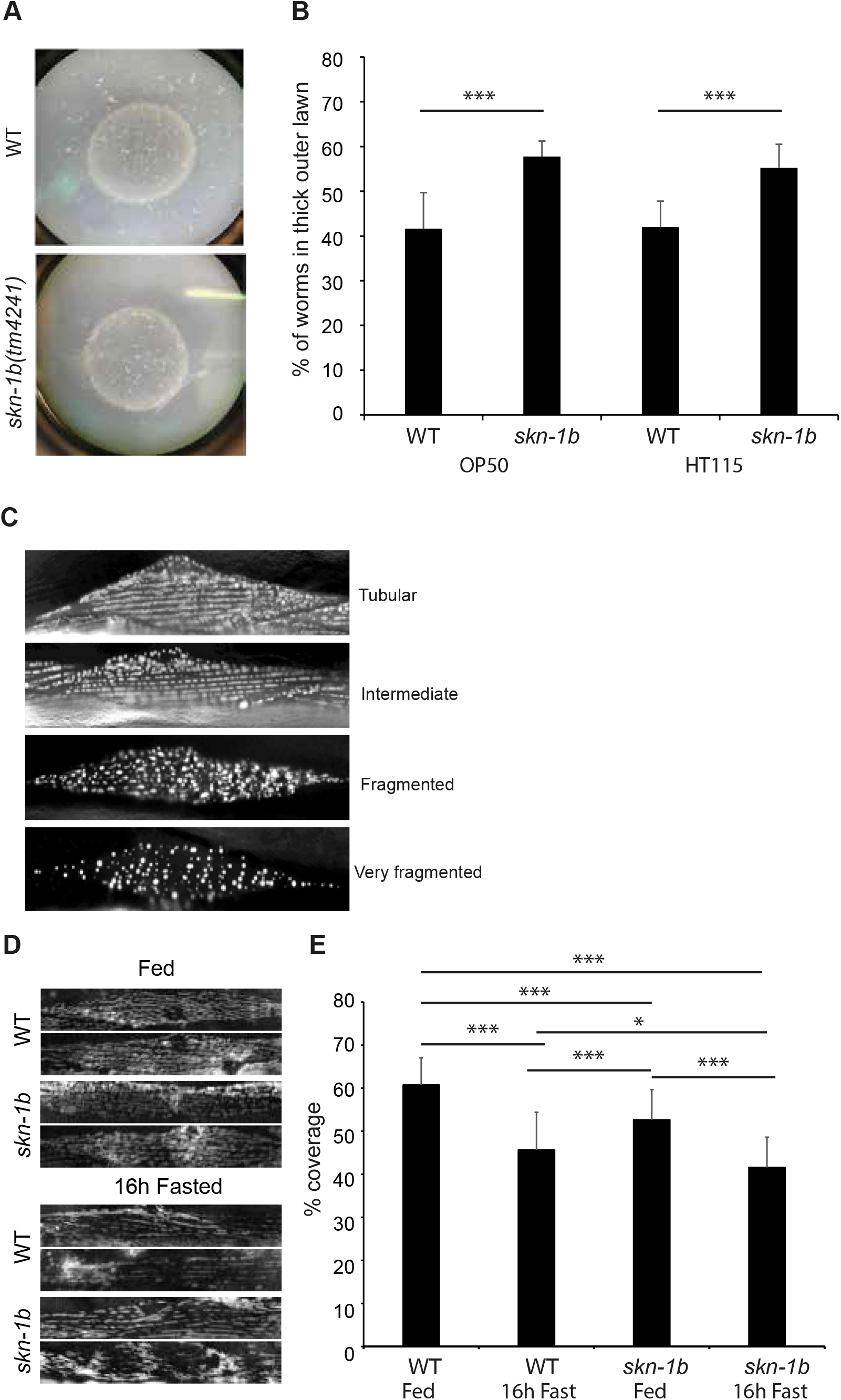
Comparing mitochondria in WT and *skn-1b* mutants - supporting data. **A and B)** Images and quantification of bordering behaviour. Each bar represents a mean of 3 biological replicates ± st. dev. **C)** Scoring system of the expression of *myo-3::mitoGFP* in *C. elegans.* **D and E)** Expression and quantification of WT and *skn-1b* mutant *C. elegans* expressing *tomm2O::GFP.* This reporter expresses a peptide of tomm20, an outer mitochondrial membrane protein and hence marks all mitochondria, deliniating their shape (Weir et al., 2017). In **E)** Each bar represents a mean of 3 biological replicates ± SEM, n>49 day 1 adults worms per group.

**Supplementary Figure 13.**
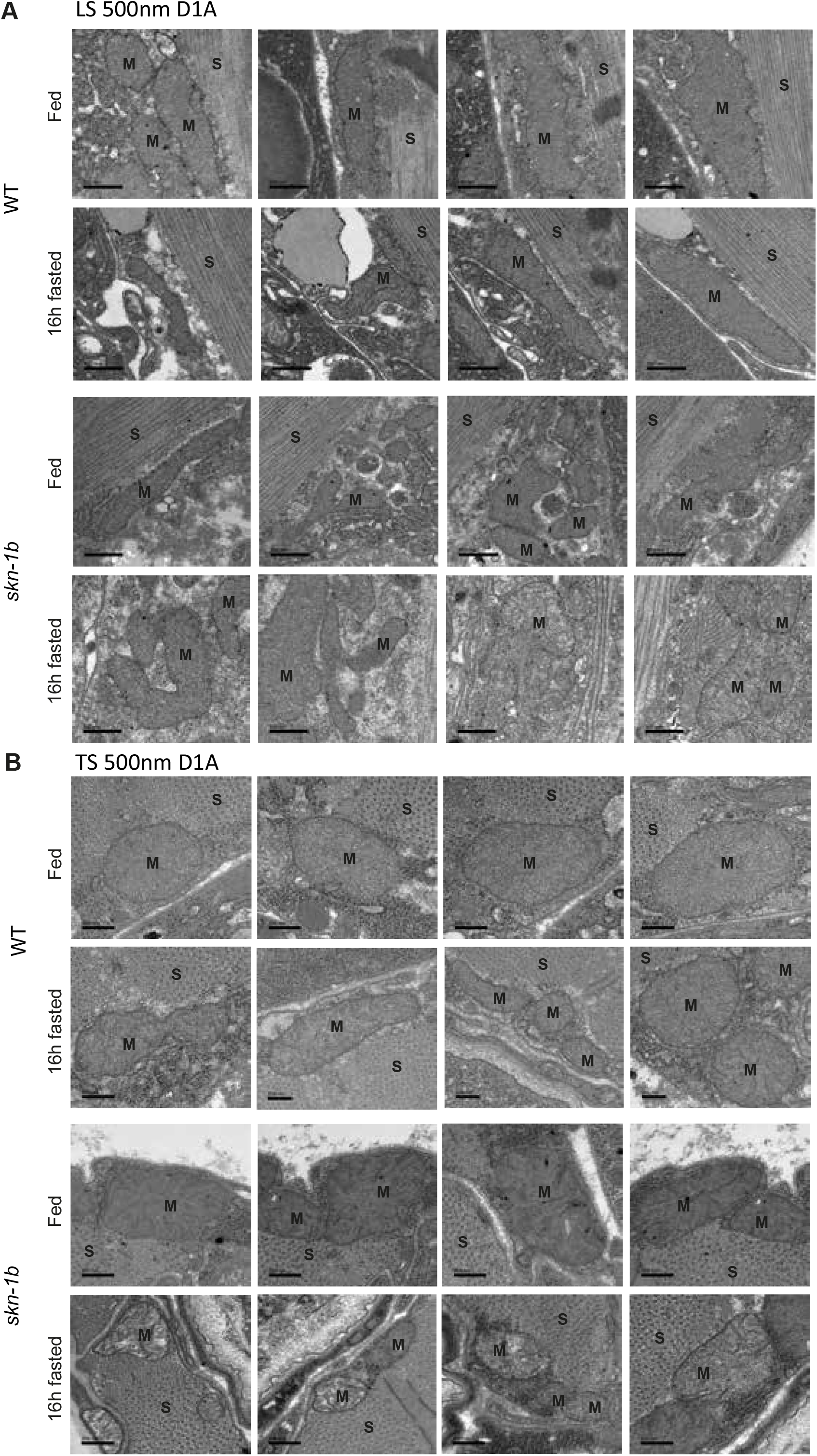
TEM of WT and *skn-1b* mutants in Fed and fasted conditions. **A)** Longitudinal sections and **B)** Transverse sections. All scale bars = 500nm, M=mitochondria, S=sarcomere. Fasting disrupts mitochondrial networks in response to fasting in WT animals. *skn-1b* mutants also have discrupted mitochondrial networks, exhibiting increased fusion of mitochondria. In reponse to fasting *skn-1b* mutant mitochondria appear much worse than WT, with disrupted membranes and critae structures.

## Notes

### Competing Interest Statement

The authors have declared no competing interest.

### Summary of Updates

Summary and introduction updated to clarify the role of SKN-1B in mediating dietary restriction incurred longevity; and author email address corrected.

## References

Alic N, Tullet JM, Niccoli T, Broughton S, Hoddinott MP, Slack C, Gems D, Partridge L. 2014. Cell-Nonautonomous Effects of dFOXO/DAF-16 in Aging. Cell Rep. doi:10.1016/j.celrep.2014.01.015

Antebi A. 2007. Genetics of aging in Caenorhabditis elegans. PLoS Genet 3:1565–1571.

Bargmann C. 2006. Chemosensation in C. elegans. WormBook. doi:10.1895/wormbook.1.123.1

Barrios A, Nurrish S, Emmons SW. 2008. Sensory Regulation of C. elegans Male Mate-Searching Behavior. Curr Biol. doi:10.1016/j.cub.2008.10.050

Ben Arous J, Laffont S, Chatenay D. 2009. Molecular and sensory basis of a food related two-state behavior in C. elegans. PLoS One. doi:10.1371/journal.pone.0007584

Birnby DA, Link EM, Vowels JJ, Tian H, Colacurcio PL, Thomas JH. 2000. A transmembrane guanylyl cyclase (DAF-11) and Hsp90 (DAF-21) regulate a common set of chemosensory behaviors in Caenorhabditis elegans. Genetics.

Bishop N a, Guarente L. 2007. Two neurons mediate diet-restriction-induced longevity in C. elegans. Nature 447:545–549. doi:10.1038/nature05904

Blackwell TK, Steinbaugh MJ, Hourihan JM, Ewald CY, Isik M. 2015. SKN-1/Nrf, stress responses, and aging in Caenorhabditis elegans. Free Radic Biol Med 1–12. doi:10.1016/j.freeradbiomed.2015.06.008

Bouret SG. 2017. Development of hypothalamic circuits that control food intake and energy balanceAppetite and Food Intake: Central Control, Second Edition. doi:10.1201/9781315120171

Bowerman B, Draper BW, Mello CC, Priess JR. 1993. The maternal gene skn-1 encodes a protein that is distributed unequally in early C. elegans embryos. Cell 74:443–452.

Bräcker LB, Siju KP, Arela N, So Y, Hang M, Hein I, Vasconcelos ML, Grunwald Kadow IC. 2013. Essential role of the mushroom body in context-dependent CO2 avoidance in drosophila. Curr Biol. doi:10.1016/j.cub.2013.05.029

Brenner S. 1974. The genetics of Caenorhabditis elegans. Genetics 77:71–94. doi:10.1002/cbic.200300625

Byrne JJ, Soh MS, Chandhok G, Vijayaraghavan T, Teoh JS, Crawford S, Cobham AE, Yapa NMB, Mirth CK, Neumann B. 2019. Disruption of mitochondrial dynamics affects behaviour and lifespan in Caenorhabditis elegans. Cell Mol Life Sci. doi:10.1007/s00018-019-03024-5

Chaudhari SN, Kipreos ET. 2017. Increased mitochondrial fusion allows the survival of older animals in diverse C. Elegans longevity pathways. Nat Commun. doi:10.1038/s41467-017-00274-4

Chávez V, Mohri-Shiomi A, Maadani A, Vega LA, Garsin DA. 2007. Oxidative stress enzymes are required for DAF-16-mediated immunity due to generation of reactive oxygen species by Caenorhabditis elegans. Genetics. doi:10.1534/genetics.107.072587

Clark LC, Hodgkin J. 2014. Commensals, probiotics and pathogens in the Caenorhabditis elegans model. Cell Microbiol. doi:10.1111/cmi.12234

Ewald CY, Landis JN, Abate JP, Murphy CT, Blackwell TK. 2014. Dauer-independent insulin/IGF-1-signalling implicates collagen remodelling in longevity. Nature 519:97–101. doi:10.1038/nature14021

Fenk LA, de Bono M. 2017. Memory of recent oxygen experience switches pheromone valence in Caenorhabditis elegans. Proc Natl Acad Sci. doi:10.1073/pnas.1618934114

Flavell SW, Pokala N, Macosko EZ, Albrecht DR, Larsch J, Bargmann CI. 2013. Serotonin and the neuropeptide PDF initiate and extend opposing behavioral states in C. Elegans. Cell 154:1023–1035. doi:10.1016/j.cell.2013.08.001

Fletcher M, Kim DH. 2017. Age-Dependent Neuroendocrine Signaling from Sensory Neurons Modulates the Effect of Dietary Restriction on Longevity of Caenorhabditis elegans. PLoS Genet. doi:10.1371/journal.pgen.1006544

Gaglia MM, Kenyon C. 2009. Stimulation of Movement in a Quiescent, Hibernation-Like Form of Caenorhabditis elegans by Dopamine Signaling. J Neurosci. doi:10.1523/jneurosci.3429-08.2009

Gallagher T, Kim J, Oldenbroek M, Kerr R, You Y-J. 2013. ASI Regulates Satiety Quiescence in C. elegans. J Neurosci. doi:10.1523/jneurosci.4493-12.2013

Ghose P, Park EC, Tabakin A, Salazar-Vasquez N, Rongo C. 2013. Anoxia-Reoxygenation Regulates Mitochondrial Dynamics through the Hypoxia Response Pathway, SKN-1/Nrf, and Stomatin-Like Protein STL-1/SLP-2. PLoS Genet. doi:10.1371/journal.pgen.1004063

Gumienny TL. 2013. TGF-β signaling in C. elegans. WormBook. doi:10.1895/wormbook.1.22.2

Hahm JH, Kim S, Paik YK. 2009. Endogenous cGMP regulates adult longevity via the insulin signaling pathway in Caenorhabditis elegans. Aging Cell. doi:10.1111/j.1474-9726.2009.00495.x

Hibshman JD, Leuthner TC, Shoben C, Mello DF, Sherwood DR, Meyer JN, Baugh LR. 2018. Nonselective autophagy reduces mitochondrial content during starvation in Caenorhabditis elegans. Am J Physiol Cell Physiol. doi:10.1152/ajpcell.00109.2018

Hoogewijs D, Houthoofd K, Matthijssens F, Vandesompele J, Vanfleteren JR. 2008. Selection and validation of a set of reliable reference genes for quantitative sod gene expression analysis in C. elegans. BMC Mol Biol 9:9. doi:10.1186/1471-2199-9-9

Hsin H, Kenyon C. 1999. Signals from the reproductive system regulate the lifespan of C. elegans. Nature 399:362–366. doi:10.1038/20694

Kapahi P, Kaeberlein M, Hansen M. 2017. Dietary restriction and lifespan: Lessons from invertebrate models. Ageing Res Rev. doi:10.1016/j.arr.2016.12.005

Kenyon CJ. 2010. The genetics of ageing. Nature 464:504–512. doi:10.1038/nature08980

Kimura KD, Riddle DL, Ruvkun G. 2011. The C. Elegans DAF-2 insulin-like receptor is abundantly expressed in the nervous system and regulated by nutritional status. Cold Spring Harb Symp Quant Biol. doi:10.1101/sqb.2011.76.010660

Kobayashi A, Tsukide T, Miyasaka T, Morita T, Mizoroki T, Saito Y, Ihara Y, Takashima A, Noguchi N, Fukamizu A, Hirotsu Y, Ohtsuji M, Katsuoka F, Yamamoto M. 2011. Central nervous system-specific deletion of transcription factor Nrf1 causes progressive motor neuronal dysfunction. Genes to Cells. doi:10.1111/j.1365-2443.2011.01522.x

Kumar S, Egan BM, Kocsisova Z, Schneider DL, Murphy JT, Diwan A, Kornfeld K. 2019. Lifespan Extension in C. elegans Caused by Bacterial Colonization of the Intestine and Subsequent Activation of an Innate Immune Response. Dev Cell. doi:10.1016/j.devcel.2019.03.010

Lakowski B, Hekimi S. 1998. The genetics of caloric restriction in Caenorhabditis elegans. Proc Natl Acad Sci U S A 95:13091–13096. doi:10.1073/pnas.95.22.13091

Lapierre LR, Hansen M. 2012. Lessons from C. elegans: Signaling pathways for longevity. Trends Endocrinol Metab. doi:10.1016/j.tem.2012.07.007

Lehrbach NJ, Ruvkun G. 2019. Endoplasmic reticulum-associated SKN-1A/Nrf1 mediates a cytoplasmic unfolded protein response and promotes longevity. Elife. doi:10.7554/eLife.44425

Lehrbach NJ, Ruvkun G. 2016. Proteasome dysfunction triggers activation of SKN-1A/Nrf1 by the aspartic protease DDI-1. Elife 5. doi:10.7554/eLife.17721.001

Libina N, Berman JR, Kenyon C. 2003. Tissue-specific activities of C. elegans DAF-16 in the regulation of lifespan. Cell 115:489–502.

Lipton J. 2004. Mate Searching in Caenorhabditis elegans: A Genetic Model for Sex Drive in a Simple Invertebrate. J Neurosci. doi:10.1523/jneurosci.1746-04.2004

McCloskey RJ, Fouad AD, Churgin MA, Fang-Yen C. 2017. Food responsiveness regulates episodic behavioral states in Caenorhabditis elegans. J Neurophysiol. doi:10.1152/jn.00555.2016

McFarland DJ. 1977. Decision making by animals. Nature. doi:citeulike-article-id:3179069

McMullan R, Anderson A, Nurrish S. 2012. Behavioral and immune responses to infection require Gαq-RhoA signaling in C. elegans. PLoS Pathog. doi:10.1371/journal.ppat.1002530

Moroz N, Carmona JJ, Anderson E, Hart AC, Sinclair D a., Blackwell TK. 2014. Dietary restriction involves NAD +-dependent mechanisms and a shift toward oxidative metabolism. Aging Cell 13:1075–1085. doi:10.1111/acel.12273

Murakami M, Koga M, Ohshima Y. 2001. DAF-7/TGF-β expression required for the normal larval development in C. elegans is controlled by a presumed guanylyl cyclase DAF-11. Mech Dev. doi:10.1016/S0925-4773(01)00507-X

Murphy CT, Lee SJ, Kenyon C. 2007. Tissue entrainment by feedback regulation of insulin gene expression in the endoderm of Caenorhabditis elegans. Proc Natl Acad Sci U S A 104:19046–19050. doi:10.1073/pnas.0709613104

Oliveira RP, Abate JP, Dilks K, Landis J, Ashraf J, Murphy CT, Blackwell TK. 2009. Condition-adapted stress and longevity gene regulation by Caenorhabditis elegans SKN-1/Nrf. Aging Cell 8:524–541. doi:10.1111/j.1474-9726.2009.00501.x

Palikaras K, Lionaki E, Tavernarakis N. 2015. Coordination of mitophagy and mitochondrial biogenesis during ageing in C. elegans. Nature. doi:10.1038/nature14300

Patterson GI, Padgett RW. 2000. TGFβ-related pathways: Roles in Caenorhabditis elegans development. Trends Genet. doi:10.1016/S0168-9525(99)01916-2

Pierce SB, Costa M, Wisotzkey R, Devadhar S, Homburger S a, Buchman a R, Ferguson KC, Heller J, Platt DM, Pasquinelli a a, Liu LX, Doberstein SK, Ruvkun G. 2001. Regulation of DAF-2 receptor signaling by human insulin and ins-1, a member of the unusually large and diverse C-elegans insulin gene family. Genes Dev 15:672–686. doi:10.1101/gad.867301

Podshivalova K, Kerr RA, Kenyon C. 2017. How a Mutation that Slows Aging Can Also Disproportionately Extend End-of-Life Decrepitude. Cell Rep. doi:10.1016/j.celrep.2017.03.062

Samuel BS, Rowedder H, Braendle C, Félix M-A, Ruvkun G. 2016. *Caenorhabditis elegans* responses to bacteria from its natural habitats. Proc Natl Acad Sci. doi:10.1073/pnas.1607183113

Schmeisser S, Priebe S, Groth M, Monajembashi S, Hemmerich P, Guthke R, Platzer M, Ristow M. 2013. Neuronal ROS signaling rather than AMPK/sirtuin-mediated energy sensing links dietary restriction to lifespan extension. Mol Metab 2:92–102. doi:10.1016/j.molmet.2013.02.002

Sebastián D, Palacín M, Zorzano A. 2017. Mitochondrial Dynamics: Coupling Mitochondrial Fitness with Healthy Aging. Trends Mol Med. doi:10.1016/j.molmed.2017.01.003

Shaw WM, Luo S, Landis J, Ashraf J, Murphy CT. 2007. The C. elegans TGF-beta Dauer pathway regulates longevity via insulin signaling. Curr Biol 17:1635–1645. doi:10.1016/j.cub.2007.08.058

Skora S, Mende F, Zimmer M. 2018. Energy Scarcity Promotes a Brain-wide Sleep State Modulated by Insulin Signaling in C. elegans. Cell Rep. doi:10.1016/j.celrep.2017.12.091

Spurlock B, Tullet JMA, Hartman JL, Mitra K. 2020. Interplay of mitochondrial fission-fusion with cell cycle regulation: Possible impacts on stem cell and organismal aging. Exp Gerontol. doi:10.1016/j.exger.2020.110919

Trojanowski NF, Raizen DM. 2016. Call it Worm Sleep. Trends Neurosci. doi:10.1016/j.tins.2015.12.005

Tullet JMA, Hertweck M, An JH, Baker J, Hwang JY, Liu S, Oliveira RP, Baumeister R, Blackwell TK. 2008. Direct Inhibition of the Longevity-Promoting Factor SKN-1 by Insulin-like Signaling in C. elegans. Cell. doi:10.1016/j.cell.2008.01.030

Van Der Klaauw AA, Farooqi IS. 2015. The hunger genes: Pathways to obesity. Cell. doi:10.1016/j.cell.2015.03.008

Wang J. 2003. Global analysis of dauer gene expression in Caenorhabditis elegans. Development. doi:10.1242/dev.00363

Weir HJ, Yao P, Huynh FK, Escoubas CC, Goncalves RL, Burkewitz K, Laboy R, Hirschey MD, Mair WB. 2017. Dietary Restriction and AMPK Increase Lifespan via Mitochondrial Network and Peroxisome Remodeling. Cell Metab. doi:10.1016/j.cmet.2017.09.024

Wilson MA, Iser WB, Son TG, Logie A, Cabral-Costa J V., Mattson MP, Camandola S. 2017. Skn-1 is required for interneuron sensory integration and foraging behavior in Caenorhabditis elegans. PLoS One. doi:10.1371/journal.pone.0176798

Wong R, Piper MDW, Wertheim B, Partridge L. 2009. Quantification of food intake in Drosophila. PLoS One. doi:10.1371/journal.pone.0006063

Wu Z, Isik M, Moroz N, Steinbaugh MJ, Zhang P, Blackwell TK. 2019. Dietary Restriction Extends Lifespan through Metabolic Regulation of Innate Immunity. Cell Metab. doi:10.1016/j.cmet.2019.02.013

Yang JS, Nam HJ, Seo M, Han SK, Choi Y, Nam HG, Lee SJ, Kim S. 2011. OASIS: Online application for the survival analysis of lifespan assays performed in aging research. PLoS One 6. doi:10.1371/journal.pone.0023525

You Y jai, Kim J, Raizen DM, Avery L. 2008. Insulin, cGMP, and TGF-β Signals Regulate Food Intake and Quiescence in C. elegans: A Model for Satiety. Cell Metab. doi:10.1016/j.cmet.2008.01.005

Zhang F, Berg M, Dierking K, Félix MA, Shapira M, Samuel BS, Schulenburg H. 2017. Caenorhabditis elegans as a model for microbiome research. Front Microbiol 8. doi:10.3389/fmicb.2017.00485

Zhang J, Holdorf AD, Walhout AJ. 2017. C. elegans and its bacterial diet as a model for systems-level understanding of host–microbiota interactions. Curr Opin Biotechnol. doi:10.1016/j.copbio.2017.01.008

Zhao Y, Gilliat AF, Ziehm M, Turmaine M, Wang H, Ezcurra M, Yang C, Phillips G, McBay D, Zhang WB, Partridge L, Pincus Z, Gems D. 2017. Two forms of death in ageing Caenorhabditis elegans. Nat Commun. doi:10.1038/ncomms15458

